# Metabolic dysregulation induces impaired lymphocyte memory formation during severe SARS-CoV-2 infection

**DOI:** 10.1101/2021.12.06.471421

**Authors:** Sanjeev Gurshaney, Anamaria Morales Alvarez, Kevin Ezhakunnel, Andrew Manalo, Thien-Huong Huynh, Nhat-Tu Le, Daniel S. Lupu, Stephen J. Gardell, Hung Nguyen

**Author notes:** Corresponding Author: Hung Nguyen, PhD, 6900 Lake Nona Blvd, Orlando, FL 32827, USA.

## Abstract

Cellular metabolic dysregulation is a consequence of COVID-19 infection that is a key determinant of disease severity. To understand the mechanisms underlying these cellular changes, we performed high-dimensional immune cell profiling of PBMCs from COVID-19-infected patients, in combination with single cell transcriptomic analysis of COVID-19 BALFs. Hypoxia, a hallmark of COVID-19 ARDS, was found to elicit a global metabolic reprogramming in effector lymphocytes. In response to oxygen and nutrient-deprived microenvironments, these cells shift from aerobic respiration to increase their dependence on anaerobic processes including glycolysis, mitophagy, and glutaminolysis to fulfill their bioenergetic demands. We also demonstrate metabolic dysregulation of ciliated lung epithelial cells is linked to significant increase of proinflammatory cytokine secretion and upregulation of HLA class 1 machinery. Augmented HLA class-1 antigen stimulation by epithelial cells leads to cellular exhaustion of metabolically dysregulated CD8 and NK cells, impairing their memory cell differentiation. Unsupervised clustering techniques revealed multiple distinct, differentially abundant CD8 and NK memory cell states that are marked by high glycolytic flux, mitochondrial dysfunction, and cellular exhaustion, further highlighting the connection between disrupted metabolism and impaired memory cell function in COVID-19. Our findings provide novel insight on how SARS-CoV-2 infection affects host immunometabolism and anti-viral response during COVID-19.

**Graphical Abstract:** 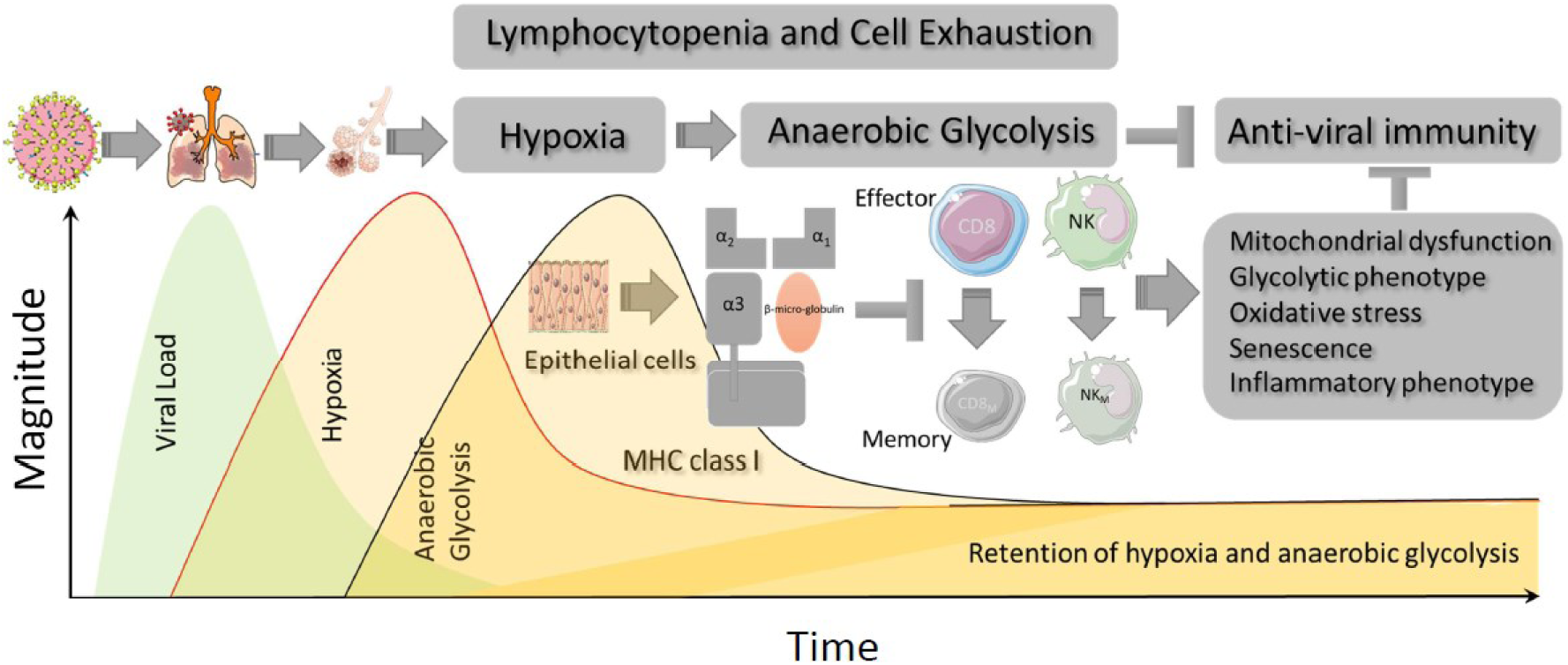

**Highlights:**

- Hypoxia and anaerobic glycolysis drive CD8, NK, NKT dysfunction
- Hypoxia and anaerobic glycolysis impair memory differentiation in CD8 and NK cells
- Hypoxia and anaerobic glycolysis cause mitochondrial dysfunction in CD8, NK, NKT cells

## Introduction

According to the World Health Organization (WHO), over 3.6 million individuals across the world have perished due to the COVID-19 pandemic as of October 2021^1^. Although multiple SARS-CoV-2 vaccines have been launched, herd immunity may not be reached in many countries until late 2021 or early 2022^2^. Additionally, mutant COVID-19 strains including the novel delta variant, which have the potential to partially evade immunity induced by currently available vaccines, as well as display significantly increased rates of infection are rapidly increasing in prevalence ^3^. Therefore, novel therapeutics to combat SARS-CoV-2 are urgently needed.

Metabolic syndrome and its cluster of conditions pose risk factors for severe COVID-19 pathogenesis^4,5^. The growing body of evidence suggests that individuals with pre-existing metabolic comorbidities are at far higher risk of suffering severe complications from COVID-19 ^6,7^. However, understanding about the metabolism of immune cells in the microenvironment of injured organs such as the lung during SARS-CoV-2 infection is limited. Most studies have been performed on patients’ peripheral blood mononuclear cells (PBMCs)^4,5,8^. Because the metabolic characteristics of the target organ and circulation are different^9^, knowledge of the immunometabolic landscape in injured organ by SARS-CoV-2 is essential for generating safe and effective treatments for COVID-19. Additionally, metabolic biomarkers, used stand-alone or in combination, that specifically predict the prognosis of COVID-19 may provide the essential knowledge for clinicians to triage care accordingly if needed.

The lung is the primary target organ of SARS-CoV-2, as the spike protein directly binds to ACE2 receptors expressed on the surface of lung epithelial cells (ECs)^10^. As a result, acute respiratory distress syndrome (ARDS) often occurs in severe COVID-19, resulting in decreased blood oxygen saturation level (hypoxia), as well as increased serum lactate dehydrogenase (LDHA) levels ^11–13^. Both downstream hypoxia signaling and hyperlactatemia have been associated with proinflammatory cytokine syndrome and lymphocyte dysfunction^14,15^. However, it is not completely understood whether/how hypoxia in COVID-19 ARDS patient affects the metabolic phenotype of immune cells. Thus, it is essential to identify metabolic dysregulation responsible for immunological impairment during severe COVID-19.

In the current study, a publicly available single cell sequencing dataset from the bronchoalveolar lavage fluid (BALF) of COVID-19 patients was used to generate a transcriptional landscape of metabolic activity at a single-cell resolution in COVID-19 lung microenvironment. We also performed high- dimensional flow cytometry of hospitalized COVID-19 patient PBMCs to validate the results at the protein level. We found that metabolic disorder by hypoxia and anaerobic glycolysis impairs memory differentiation in CD8 and NK cells during SARS-CoV-2 infection. Multiple distinct, highly resolved CD8 and NK cell subsets with metabolic and mitochondrial dysfunction were found to effectively indicate COVID-19 severity. The findings in current study provide a single-cell metabolic landscape of COVID-19, highlighting the metabolic plasticity and heterogeneity of immune cells, which linked to host immunity against SARS-CoV-2 thereby having translational application for COVID-19 severity assessment, treatment, and therapy.

## Results

### High-dimensional flow cytometry and BALF transcriptomic analysis reveal cellular dysfunction and impaired memory formation in CD8 T and NK cells in patients with COVID- 19

Immunophenotyping of PBMCs from hospitalized COVID-19 patients was performed to probe the immunological response and metabolic perturbations that arise from SARS-CoV-2 infection (**Fig. 1A**). PBMCs were freshly isolated from 20 COVID-19 patients and 8 control subjects and analyzed without stimulation or cryogenic preservation (**Fig. 1A**). Given their critical roles in the primary immune response against SARS-CoV-2, we initially start with CD8^+^T and NK cells^16,17^. Principal component analysis (PCA) revealed distinct clustering of COVID-19 and healthy control samples, thus revealing that PBMCs from COVID-19 patients exhibited an abnormal immunophenotype (**Fig. 1B**). FlowSOM unsupervised clustering and universal manifold approximation projection (UMAP) dimensionality reduction were next performed to detect unsupervised clusters. UMAP embedding confirmed distinguished immunophenotype of PBMC from COVID-19 patient and healthy control (**Fig. 1C**). Unsupervised clusters were annotated based upon canonical marker expression (Supplementary Fig. S3A-B) to define distinct and highly resolved cell populations (**Fig. 1D, E**). Cytotoxic T lymphocytes (CTLs) were identified by co-expression of CD8 and GZMB; CD8 central memory by CD8, CD62L, and CCR7; CD8 transitional memory by CD8, CD62L; CD8 effector memory by CD8, CCR7, GZMB; NK by CD56; NKT by CD56, CD8; memory NK (CD62L^+^NK) by CD56, CD62L (Supplementary Table 3).

**Fig. 1.**
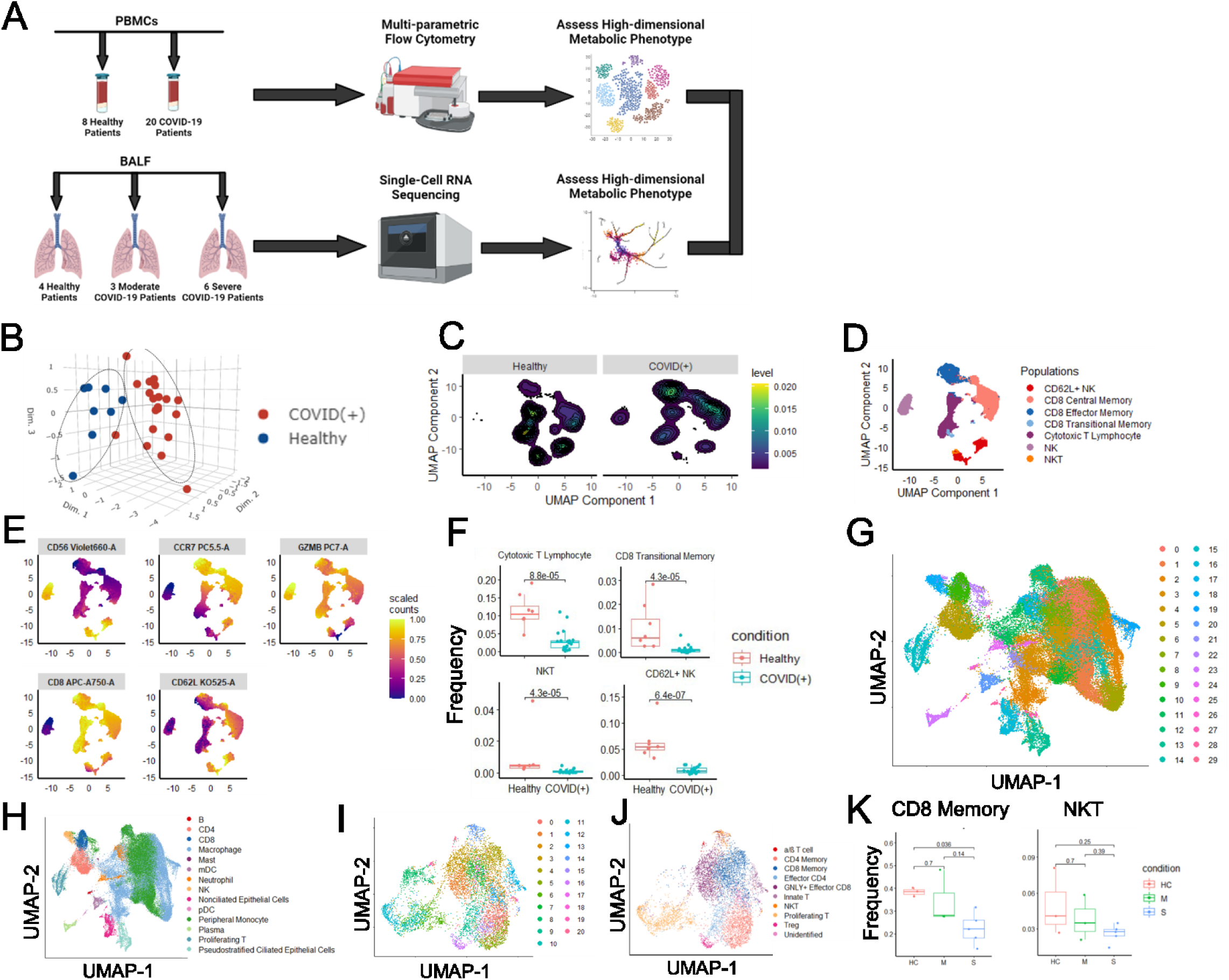
Distinct immunophenotype of BALFs and PBPMCs from COVID-19 patients. **A.** Schematic illustrating experimental design for multiparametric flosw cytometry and single-cell RNA sequencing re-analysis. High-dimensional flow cytometry of freshly isolated PBMCs from 8 Healthy Control and 20 hospitalized COIVD-19 patients (B-F). **B.** 3-D PCA analysis conducted using bulk expression of each marker per sample as input, circles were manually drawn around the PCA plot to highlight distinct clustering. **C.** Contour plot of kernel density for UMAP projection of PBMCs, 5,000 cells from each condition (Healthy and COVID(+)) were randomly subsetted for density analysis**; D.** UMAP projection of labelled PBMC populations from Healthy (8) and COVID- 19 (20) patients; **E.** UMAP projections of PBMCs overlayed with scaled expression of markers used for unsupervised clustering; **F.** Box-plot of cell-type proportion for each disease state, dot represents individual sample, 2-sided Wilcoxon Mann Whitney test was performed to indicate statistical significance; Immunophenotyping of scRNA-seq data of isolated BALFs derived from healthy control, moderate and severe COVID-19 patients (F-J); **G.** UMAP projection displaying unsupervised clusters of 66,452 cells from Healthy Control (4), Moderate (3), Severe (6) patients; **H.** UMAP projection of labelled BALF populations annotated based upon canonical marker expression; **I.** UMAP projection displaying unsupervised clusters of 7601 reintegrated T cells from Healthy Control (3), Moderate (3), Severe (5); **J.** UMAP projection of labelled reintegrated T cells annotated based upon canonical marker expression; **K.** Box-plot of cell-type proportion of CD8 Memory and NKT for each disease state, dot represents individual sample, 2-sided Wilcoxon Mann Whitney test was performed to indicate statistical significance.

Nonparametric differential abundance testing showed significantly decreased levels of CD8 transitional memory and memory NK cells in PBMCs from COVID-19 patients (**Fig. 1F**) Additionally, the percentages of CTLs and NKT were remarkably reduced in COVID-19 PBMCs. These findings suggest that severe COVID-19 infection may be associated with impaired lymphocyte memory formation and accompanying immunological dysfunction.

We also examined if these changes were also identified in cells harvested from the lungs of COVID-19 patients, the primary target for virus infection, using publicly available single cell RNA-sequencing data from BALF samples^18^. Louvain optimization and UMAP dimensionality reduction were applied to generate unsupervised clusters (**Fig. 1G**) which were then annotated by expression of canonical markers (Supplementary Fig. 1A, B) to define highly resolved populations (**Fig. 1H**). Canonical markers used to annotate the unsupervised clusters and define cell subsets were as follows: CD8^+^ T cells (*CD3D* and *CD8A)*; CD4^+^T cells (*CD3D* and *CD4)*; proliferating T cells (*CD3D* and *MKI67)*; pseudostratified epithelial cells (*EPCAM*, *CFAP126*, and *DNAAF1)*; non-ciliated epithelial cells (*EPCAM)*; plasma cells (*IGKC* and *MS4A1*(-)); B cells (*MS4A1)*; neutrophils (*HCAR3)*; NK cells (*KLRC1)*; MAST cells (*LTC4S)*; pDCs (*CLEC4C)*; myeloid dendritic cells (mDCs) (*CD1C* and *CLEC9A)*; macrophages (*CD68 and FABP4)*; peripheral monocytes (*CD68*, *FCN1*, and *CD14)* (Supplementary Fig. 1A-B).

After initial cell type identification, T cells were subsetted and unsupervised clustering was performed again to achieve higher resolution into distinct T cell lineages (**Fig. 1I, J** and Supplementary Fig. 2A). Abundance of CD8 memory and NKT cells was found reduced and positively correlated to disease severity (**Fig. 1K**). Taken together, these results suggest that memory differentiation of CD8^+^T and NK cells is impaired in both the lungs and in circulation of COVID-19 patients.

### Hypoxia/anaerobic glycolysis axis causes cellular dysfunction of CTLs in COVID-19

Expression of cellular markers for metabolism and exhaustion was evaluated in CD8 T cells from COVID-19^+^ hospitalized patients to gain insight about the mechanistic underpinnings of cellular dysfunction. Increased *HIF-1α* expression in CD8 T cells is indicative of hypoxia, a condition that accompanies pulmonary damage in COVID-19 patients (Supplementary Fig. 4A). Increased glycolytic dependence of CD8 T cells from COVID-19 patients was suggested by the upregulation of 2-(2-(N-(7-Nitrobenz-2-oxa-1,3-diazol-4-yl)Amino)-2-deoxyglucose) (2-NBDG) uptake (Supplementary Fig. 4B). The glucose transporter 1 (GLUT1) has been well known to govern glycolytic flux ^19^ and participates in the generation of cytotoxic CD8 lymphocytes (CTLs)^20^. The frequency and expression of GLUT1 in CTLs present in PBMCs from COVID-19 patients were significantly increased (Supplementary Fig. 4C, D), suggesting that these cells are phenotypically glycolytic. Under hypoxic conditions, a metabolic phenotype primarily dependent on anaerobic glycolysis often leads to oxidative stress and mitochondrial dysfunction^21^. Indeed, the frequency of ROS^+^CTL was elevated in PBMCs from COVID-19 patients (Supplementary Fig. 4E). Furthermore, GLUT1^+^CTL of COVID-19 patients expressed higher levels of VDAC-1 (Supplementary Fig. 4F). which is an indicator of reactive oxidative species (ROS) generation and mitochondrial death^22^. This metabolic dysregulation in COVID-19 CTLs is likely linked to cellular exhaustion, as indicated by the upregulation of LAG-3 in this cell subset. (Supplementary Fig. 4G). To further establish the interplay between glycolysis, mitochondrial dysfunction, and cellular exhaustion in CTLs, we performed unsupervised clustering specifically on metabolic and functional markers to glean insight into the metabolic state of CTLs (Supplementary Fig. 3C). We observed and defined a unique cluster displaying high expression of GLUT1, LAG-3, ROS, and VDAC1, indicative of elevated glycolysis, exhaustion, and mitochondrial stress, as GLUT1^+^ mitochondrially exhausted CTLs (Supplementary Fig. 4H). This cell population was enriched in severe COVID-19 PBMCs (Supplementary Fig. 4I). These results indicate that mitochondrial and metabolic dysfunction that is closely tied to CTL exhaustion in COVID-19 patients.

The BALF transcriptomic dataset also revealed increased expression of genes encoding anaerobic glycolysis in CTLs from moderate (M)- and severe (S)- COVID-19 patients (Supplementary Fig. 5A,B). Expression of transcripts coding for glycolytic enzymes including *GAPDH, GALM,* and *ALDOA* was significantly increased in COVID-19 CTLs (Supplementary Fig. 5A,B). Key metabolic pathways including hypoxia, anaerobic glycolysis, mitophagy, autophagy, cell exhaustion, and senescence were upregulated, while pathways relying on mitochondrial metabolism including fatty acid oxidation (FAO), cholesterol metabolism, and OXPHOS were attenuated in CTLs from COVID-19 patients (Supplementary Fig. 5A,C). The increased glycolytic state in S-COVID-19 CTLs was revealed by GSEA analysis coupled with the overexpression of key glycolytic regulator genes (*GAPDH, GALM, ALDOA)* (Supplementary Fig. 5A,B). Hierarchical clustering of differentially expressed glycolytic genes suggested a tight association between *HIF- 1α* expression and anaerobic glycolysis (Supplementary Fig. 5B), indicating that oxygen-deprived condition in the BALF environment links to glycolytic metabolism. During normal CTL differentiation, cells preferentially shift from OXPHOS towards mTOR/ATK-mediated aerobic respiration to sustain increased bioenergetics demand and augmented mitochondrial biosynthesis^23^. GSEA analysis showed that CTLs are less dependent on mitochondrial TCA metabolism when impacted by SARS-CoV-2 infection (Supplementary Fig. 5C). Reduction of NAD^+^ to NADH conversion is required to preserve cellular redox homeostasis and sustain glycolytic flux. We observed decreased expression of transcripts encoding NADH oxidoreductases (*NDUFB8, NDUFC2, and NDUFA11)* in CTLs from COVID-19 patients (Supplementary Fig. 5A,C). There was also downregulation of lipid metabolism- associated genes (*FABP4, APOC1, APOE, MARCO*) in COVID-19 CTLs (Supplementary Fig. 5A,C). Increased oxidative stress in COVID-19 CTLs was evident by overexpression of *NFE2L2* and *PRDX2* (Supplementary Fig. 5D). Decreased NADH oxidation and a concomitant increased NAD^+^ level is associated with impaired cytokine secretion, cell proliferation, and exhaustion^24,25^. Indeed, the expression of *CD38*, an NAD^+^ hydrolase linked to T cell exhaustion ^25^, was increased in COVID- 19 CTLs (Supplementary Fig. 5A,B). These data reveal that hypoxia- induced CD38 expression is associated with metabolic reprograming and cellular exhaustion of CTLs in the lung of COVID- 19 patients. This conclusion is supported by higher levels of exhaustion marker *LAG3* in CTLs from S- COVID-19 patients (Supplementary Fig. 5A,B). Taken together, these results suggest that hypoxia arising from COVID-19-pulmonary dysfunction triggers impaired FA metabolism and oxidative stress, which induces mitochondrial dysfunction and cellular exhaustion.

### Metabolic dysregulation in COVID-19 impaired CD8 T-cell memory differentiation

Memory cell differentiation of CD8 T cells is a critical component of the immunological response against viral re-infection^26^. To better understand the kinetics and dynamics of CD8 memory cell differentiation during SARS-CoV-2 infection, we performed trajectory inference and pseudotemporal modeling analysis on BALF CD8 T cells (**Fig. 2A**). Differential analysis revealed decreased pseudotime values for CD8 memory in S compared to M- COVID-19 and healthy control (**Figs. 2A, B)**. The frequencies of proliferating CD8 and GNLY^+^ effector CD8 T cells were significantly increased in severe COVID-19 compared to healthy controls (**Fig. 2C**). Moreover, CD8 cells were found highly proliferating in M- than S- COVID-19 patients (**Fig. 2C**). In contrast, a lower abundance of GNLY^+^ effector CD8 cells was found in S-COVID-19. These findings suggest that CD8 T cells are stalled along the memory differentiation trajectory and are unable to reach the terminal state during S- COVID-19 infection.

**Fig. 2.**
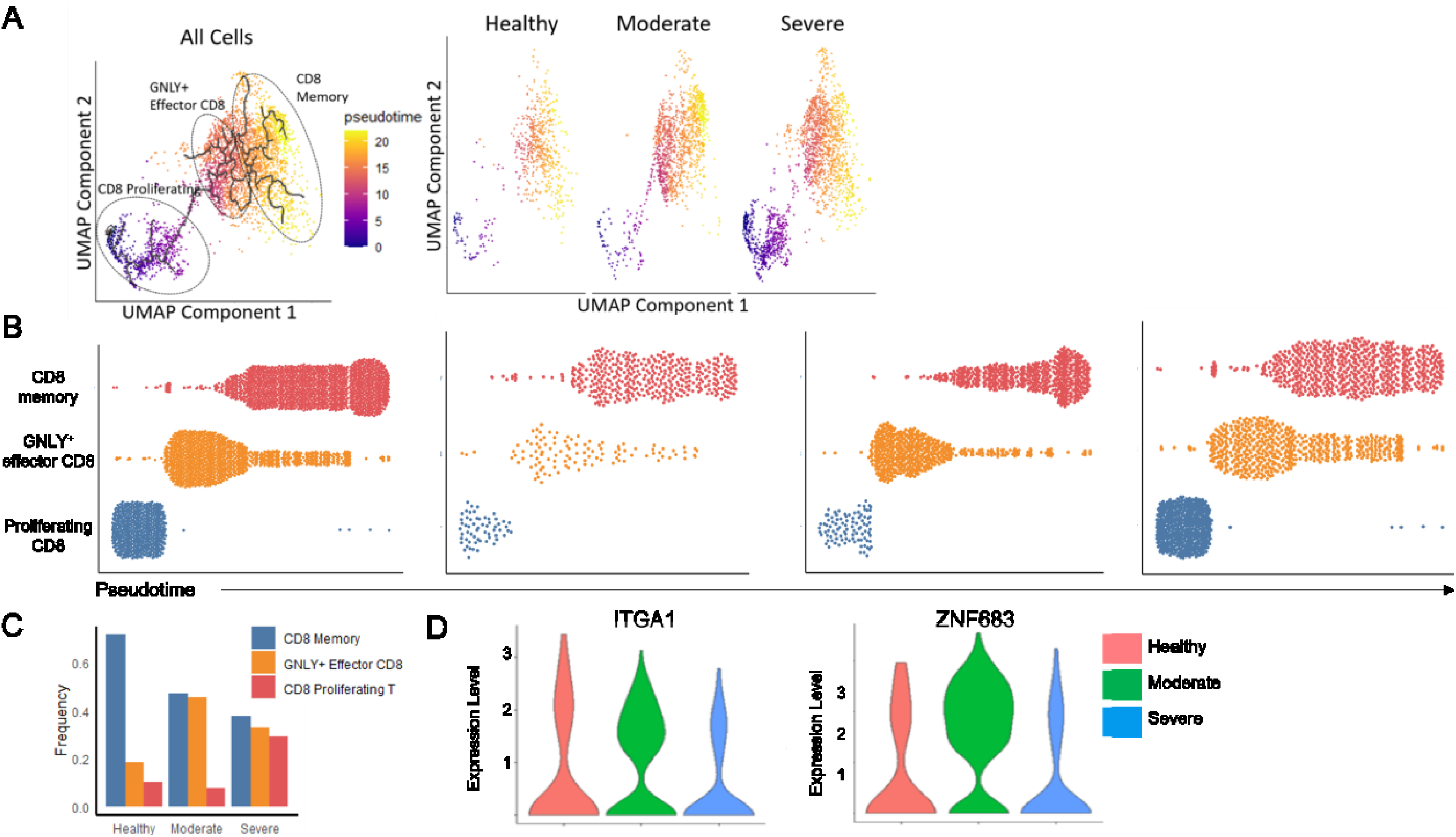
Impaired memory differentiation of CD8 T-cells in COVID-19. Pseudotime and trajectory inference analysis was conducted to evaluate the differentiation kinetics of CD8 T-cells in the BALF during SARS-CoV-2 infection. **A**. UMAP projection of 3,694 CD8 cells from all reintegrated samples, healthy samples alone, moderate samples alone, and severe samples alone, with trajectory mappings colored by pseudotime; **B**. Dot plot showing pseudotime values for CD8 cells from all reintegrated samples, healthy samples alone, moderate samples alone, and severe samples alone, each dot represents a cell; **C**. Bar graph displaying frequency of CD8 subpopulations across disease conditions; **D**. Violin plot of expression of tissue resident memory phenotype genes (*ITGA1*, *ZNF683*) in CD8 memory cells compared across disease states.

During viral infection, circulating effector memory cells migrate to the infected tissue and differentiate into tissue-resident memory (T_RM_) cells to provide the first response defense against reencounter of the pathogen ^27^. Consistently, CD8 effector memory (CD8_EM_) cells expressed higher tissue residence phenotype (i.e., increased expression of *ITGA1 and ZNF683*) in M- as compared to S- COVID-19 patients (**Fig. 2D**). This finding may indicate that impaired CD8 memory differentiation is driving factor resulting in impaired viral clearance during severe COVID-19. In contrast, trajectory inference and pseudotemporal ordering revealed no significant difference between the differentiations of CD4 cells amongst conditions (Supplementary Figs. 7A, B), suggesting that CD4 T cells retain proper differentiation and memory formation despite the metabolic stress during SARS-CoV-2 infection.

We next analyzed the metabolic landscape of CD8_M_ cells in the BALF environment. Effector BALF CD8 T cells were identified as memory cells based on expression of *HOPX and SELL* (Supplementary Fig. 3A). GSEA analysis revealed that CD8_M_ cells from S- or M- COVID-19 patients were highly dependent on glycolysis for their energetic demands (**Figs. 3A, B**). Close clustering and shared upregulation of glycolytic enzyme coding genes *GALM, GAPDH, GPI, and ALDOA* with *HIF1A*, and transcripts regulating exhaustion (*TIGIT* and *LAG3*) (**Fig. 3B**), indicated that hypoxia and anaerobic glycolysis are associated with impaired CD8_M_ function in COVID-19 BALF. FAO oxidation and OXPHOS promote the development of CD8_M_ cells after antigen exposure^28^. Indeed, genes encoding regulators of lipid uptake (*APOE,* and *APOC1*) and FAO (*OLR, MARCO, FABP4, PPRγ*) were downregulated in CD8_M_ cells of S-COVID-19 patients (**Figs. 3A, C**). Decreased expression of OXPHOS coding genes was found in S-COVID-19 CD8_M_ cells (**Figs. 3A, C**). FAO coding transcripts were negatively correlated with *HIF1A* expression (**Fig. 3C**), indicating that impaired mitochondrial metabolism may be a result of hypoxic conditions. Pearson correlation analysis revealed a negative correlation (*R = -0.73*, *p = 0.011*) between module scores for glycolysis and FAO (**Fig. 3D**), which further showed a potential association between prolonged anaerobic glycolysis and reduced mitochondrial fitness. Moreover, a strong positive correlation between module scores for glycolysis and exhaustion (*R = 0.85*, *p* = 0.00026) (**Fig. 3D**) validates that excessive glycolytic dependence is likely tightly linked to CD8_M_ exhaustion. Genes involved in cellular senescence and mitophagy were also upregulated in S-COVID-19 CD8_M_ cells (**Fig. 3A**), implying that CD8_M_ cells metabolically switch to these pathways to satisfy bioenergetics demands in response to impaired mitochondrial metabolism. Likewise, glutaminolysis is also used as an alternative bioenergetic pathway, evident by upregulation of glutamate oxidation regulating genes *GLUD1* and *DGLUCY* in CD8_M_ cells during COVID-19, likely in response to reduced lipid uptake and FAO (**Fig. 3A**).

**Fig. 3.**
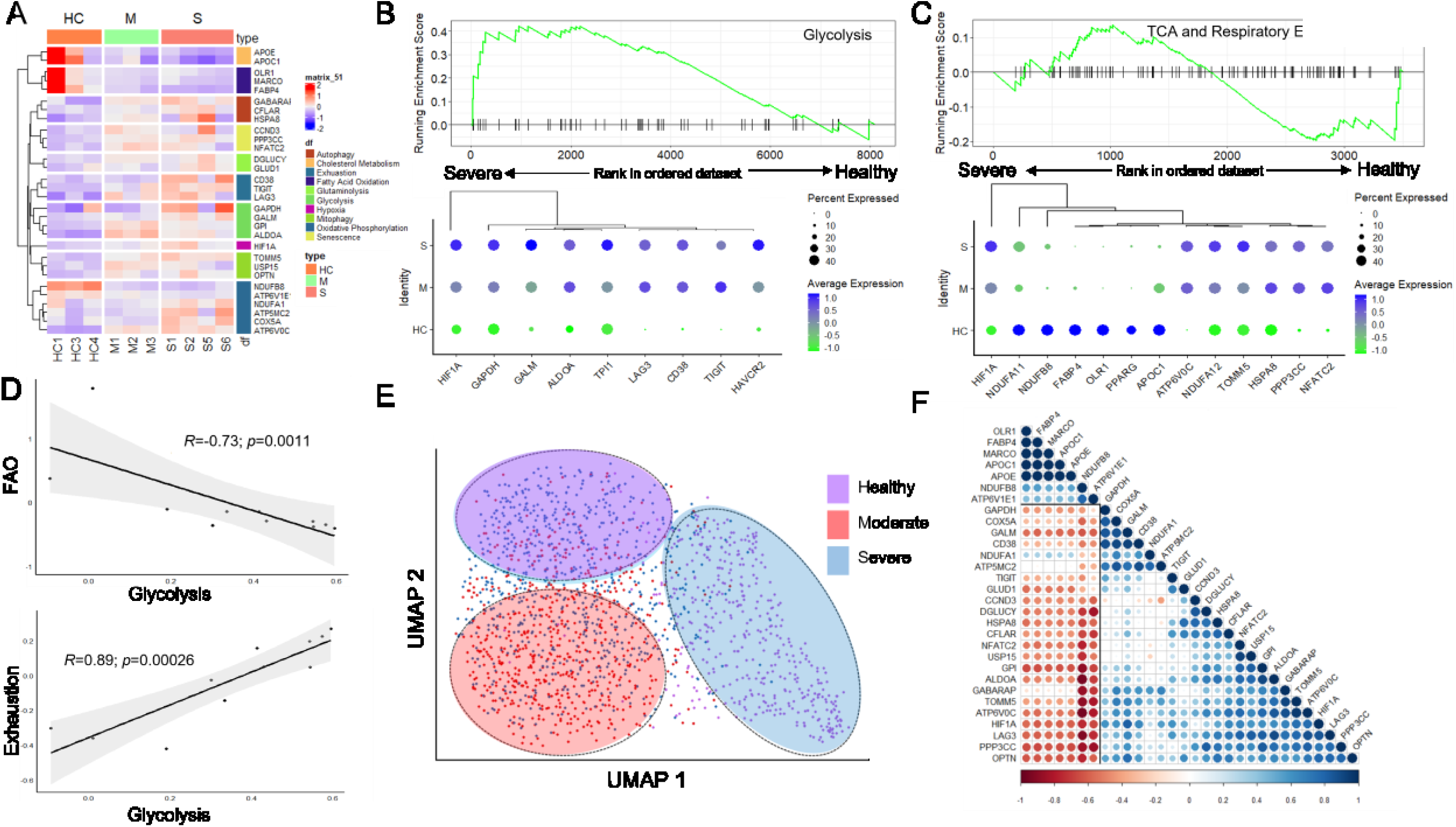
CD8 Memory Cells in the BALF undergo anaerobic metabolic reprogramming. Analysis of metabolic phenotype of CD8 memory cells from the BALF of healthy and severe COVID-19 patients; **A.** Heatmap displaying expression of key differentially expressed metabolic genes for CD8 memory cells; **B.** GSEA enrichment plots for “Glycolysis” comparing severe vs healthy control patients, adjacent is a dotplot demonstrating expression and hierarchical clustering of select key glycolytic genes; **C.** GSEA enrichment plots for “TCA and Respiratory Electron Transport” pathways comparing severe vs healthy control patients, adjacent is a dotplot demonstrating expression and hierarchical clustering of select key genes involved in mitochondrial metabolism; **D.** Linear regression and Pearson correlation analysis between module scores for Glycolysis and Exhaustion, and module scores for glycolysis and fatty acid oxidation; **E.** UMAP projections of CD8 memory cells clustered solely on the expression of 42 differentially expressed metabolic genes; **F.** Correlation matrix showing pearson correlation between differentially expressed metabolic genes

PCA analysis performed on 30 differentially expressed metabolic genes (**Table S1**) showed distinct clustering of CD8_M_ cells across different groups, further highlighting the potential of these pathways to be used as predictive markers for disease severity (**Fig. 3E**). In this line, Pearson correlation analysis showed a strong positive correlation between expression of genes regulating glycolysis, mitophagy, senescence, and glutaminolysis (**Fig. 3F**). These genes were inversely correlated with transcripts regulating FAO and NADH oxidation (**Fig. 3F**). Finally, increased CD38 expression in S- COVID-19 CD8_M_ cells was closely clustered with exhaustion-coding genes (LAG- 3 and TIGIT) (**Fig. 3B**) suggesting CD38 expression is associated with metabolic reprograming, memory impairment and cellular exhaustion of CD8_M_ in the lung of COVID-19 patients.

Because CD8 memory cell differentiation is highly dependent on their metabolism^29^, we assessed the metabolic profile of CD8_M_ cells from patient-derived PBMCs. Increased abundance of GLUT1^+^CD8_M_ cells in COVID-19 PBMCs validated the glycolytic phenotype of CD8_M_ cells in COVID-19 (**Fig. 4A**). GLUT1^+^CD8_M_ cells exhibited increased ROS expression (**Fig. 4B**), which might reflect elevated oxidative stress in COVID-19 PBMCs. Upregulation of LAG-3 in COVID-19 GLUT1^+^CD8_M_ cells was indicative of cellular exhaustion in addition to memory cell impairment in COVID-19 CD8_M_ cells (**Fig. 4C**). Despite no apparent change in ROS^+^ CD8_M_ cell abundance between diseases states (**Fig. 4D**), these cells indeed exhibited augmented HIF-1α and VDAC expression, suggesting hypoxia-induced mitochondrial dysfunction in this cell subset during COVID-19 infection (**Fig. 4E,F**). Downstream unsupervised clustering on metabolic markers revealed increased abundance of GLUT1^+^ mitochondrially exhausted CD8_M_ in COVID-19 PBMCs (**Fig. 4G**). Together, these data demonstrate that the hypoxia/anaerobic glycolysis axis mediates CD8_M_ cellular dysfunction and exhaustion in COVID-19.

**Fig. 4.**
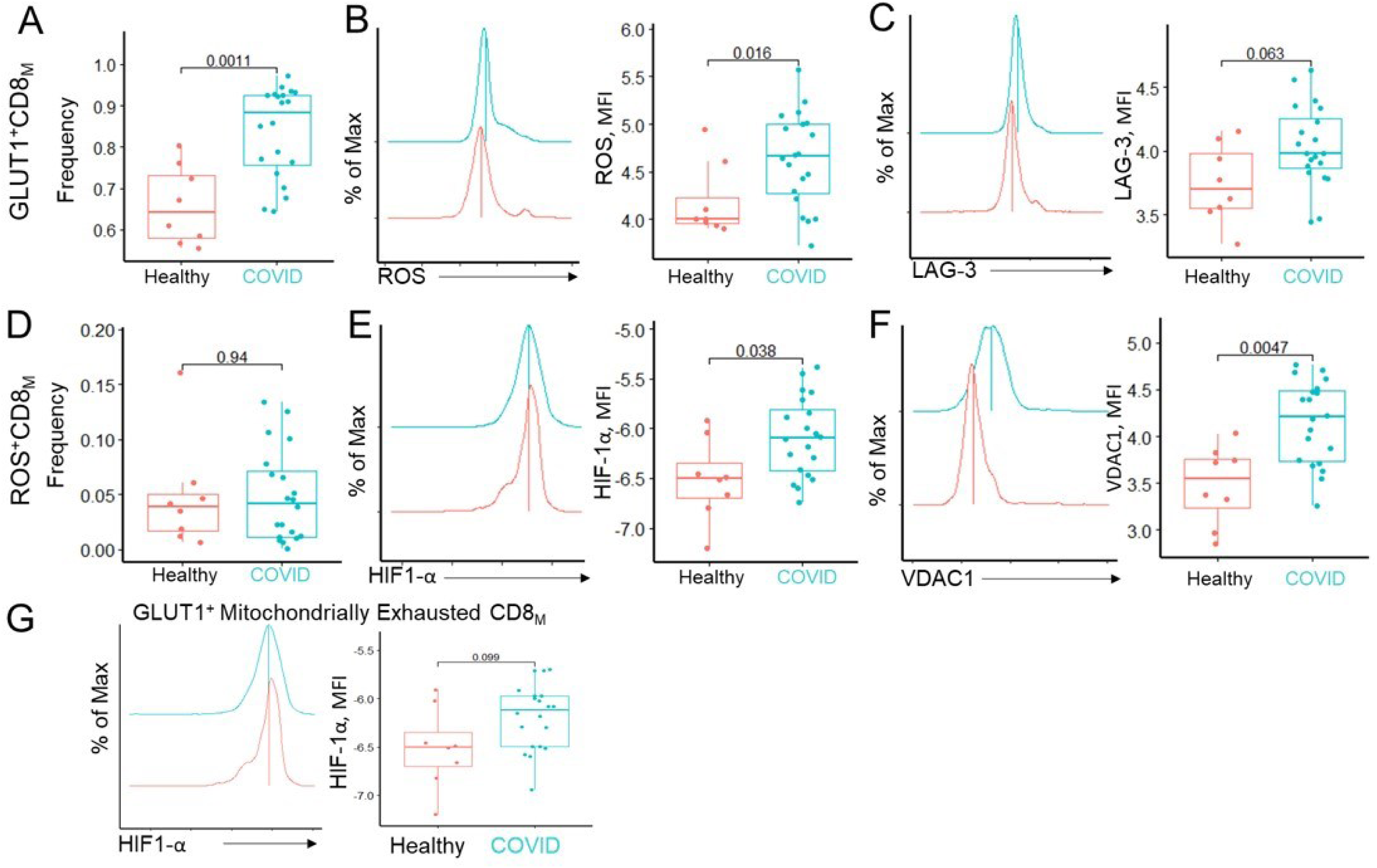
Circulating CD8 memory cells exhibit metabolic driven exhausted phenotype in COVID-19. Metabolic state and cell function of CD8 memory cells from Healthy and COVID-19 PBMCs were evaluated. **A.** Box-plot of cell-type proportion of GLUT1+ CD8 memory cells for each disease state, each dot represents individual sample, 2-sided Wilcoxon Mann Whitney test was performed to indicate statistical significance; **B.** Density plot displaying distribution of ROS expression in GLUT1+ CD8 memory cells, adjacent **to** boxplot displaying MFI values of ROS expression in GLUT1+ CD8 memory cells for both Healthy and COVID-19 patients; **C.** Density plot displaying distribution of LAG-3 expression in GLUT1+ CD8 memory cells, adjacent **to** boxplot displaying MFI values of LAG-3 expression in GLUT1+ CD8 memory cells for both Healthy and COVID-19 patients; **D.** Box-plot of cell-type proportion of ROS+ CD8 memory cells for each disease state; **E.** Density plot displaying distribution of HIF-1a expression in ROS+ CD8 memory cells, adjacent **to** boxplot displaying MFI values of HIF-1a expression in ROS+ CD8 memory cells for both Healthy and COVID-19 patients; **F.** Density plot displaying distribution of VDAC1 expression in ROS+ CD8 memory cells, adjacent **to** boxplot displaying MFI values of VDAC1 expression in ROS+ CD8 memory cells for both Healthy and COVID-19 patients; **G.** Density plot displaying distribution of HIF-1A expression in GLUT1+ Mitochondrially Exhausted CD8 memory cells, adjacent **to** boxplot displaying MFI values of HIF-1A expression in GLUT1+ Mitochondrially Exhausted CD8 memory cells for both Healthy and COVID-19 patients

### Aberrant metabolism causes NKT dysfunction in COVID-19 patients

NKT cells which express CD56 and CD8 are intermediate between the CD8 and NK cell lineages. NKT cells play critical roles in preventing pneumonia during chronic pulmonary disease^30^. There were reduced levels of circulating and BALF NKT cells in severe COVID-19 patients (**Figs. 1F**). We examined the potential link between aberrant cellular metabolism and NKT lymphocytopenia in COVID-19. The frequency of glycolysis dependent GLUT1^+^ NKT cells are more abundant in COVID-19 patients (Supplementary Fig. 8A). There was also a striking increase in the frequency of ROS^+^ NKT cells in COVID-19 PBMCs (Supplementary Fig. 8B). Interestingly, these cells displayed elevated HIF-1α expression (Supplementary Fig. 8B), which further implied that hypoxia also triggers oxidative stress in NKT cells during COVID-19 infection. To further establish the relationship between this altered metabolism and cellular function, a secondary unsupervised clustering was performed. We identified a population of GLUT1^+^ mitochondrially exhausted NKT cells, representative of combined augmented glycolytic phenotype, impaired mitochondrial function, and cellular exhaustion, that were significantly increased in COVID-19 patients (Supplementary Fig. 8C). Higher HIF-1α expression in this cell subset further validated that hypoxia-induced glycolysis is a key mechanism underlying NKT mitochondrial dysfunction (Supplementary Fig. 8C). In summary, these results suggest that NKT cells in circulation acquire significant metabolic-induced cell dysfunction because of prolonged exposure to hypoxic conditions in COVID-19.

To further probe the perturbed metabolism in NKT cells from COVID-19 patients, we characterized gene expression profiles of BALF NKT. Co-expression of *CD8A and KLRD1* was used to define NKT cell lineage (Supplementary Fig. 3A). A transcriptional program associated with hypoxia- induced metabolic reprogramming was seen in NKT cells from S COVID-19 patients (Supplementary Fig. 8D). Normally, activated NKT cells use pyruvate dehydrogenase (*PDHA1/2*) to supply acetyl-CoA to the TCA cycle^31^. However, the oxygen- deprived environment in the lung of COVID-19 patients was found to promote downregulation of genes encoding NADH oxidation (*NDUFB8, NDUFA11, NDUFA13*), as well as FAO and lipid uptake (Supplementary Fig. 8D,E). Consequently, NKT cells are dependent on anaerobic glycolytic metabolism to fulfil their bioenergetic demands (Supplementary Fig. 8D,F). However, expression of genes- regulating OXPHOS and TCA cycle were upregulated, suggesting that mitochondrial fitness was likely not affected in COVID-19 NKT cells (Supplementary Fig. 8D,E). This observation further revealed that OXPHOS was insufficient to support NKT effector function under COVID-19 hypoxic conditions. Alternatively, increased amounts of *GADD45B* and *SLC25A5* transcripts (Supplementary Fig. 8E) indicated that NKT cells undergo metabolic adaptation via enhancement of mitophagic activity to produce basal catabolic intermediates required for effector cytokine secretion.

### Dysregulated metabolism promotes memory CD62L^+^NK cell exhaustion in COVID-19 patients

CD62L^+^NK cells display memory-like attributes, evident by their rapid response to secondary viral infection and subsequent transformation to effector subtypes^32^. Decreased percentages of CD62L^+^ NK cells in COVID-19 PBMCs (**Fig. 5A**) prompted us to investigate the impact of cellular metabolism on CD62L^+^NK survival and function. The frequency of GLUT1^+^CD62L^+^NK was increased in COVID-19 PBMCs (**Fig. 5B**). Importantly, GLUT1^+^ CD62L^+^ NK frequency was positively correlated with serum glucose levels (*R = 0.76*, *p = 0.0042*) (**Fig. 5C**), which unveils a pivotal relationship between blood glucose levels and memory cell glucose uptake. GLUT1^+^ CD62L^+^ NK also expresses significantly higher levels of HIF-1α (**Fig. 5D**) and ROS (**Fig. 5E**), suggesting that hypoxia- mediating anaerobic glycolysis induces oxidative stress in CD62L^+^NK cells during COVID-19 infection. This interpretation was supported by upregulation of HIF-1α in ROS^+^CD62L^+^ NK cells (**Fig. 5F**). Given that elevated GLUT1 expression is associated with lymphocyte exhaustion and mitochondrial dysfunction, these results may suggest that hyperglycemic COVID-19 infected patients will likely exhibit exacerbated CD62L^+^ NK cell dysfunction because of hypoxia-driven glycolytic metabolic reprogramming.

**Fig. 5.**
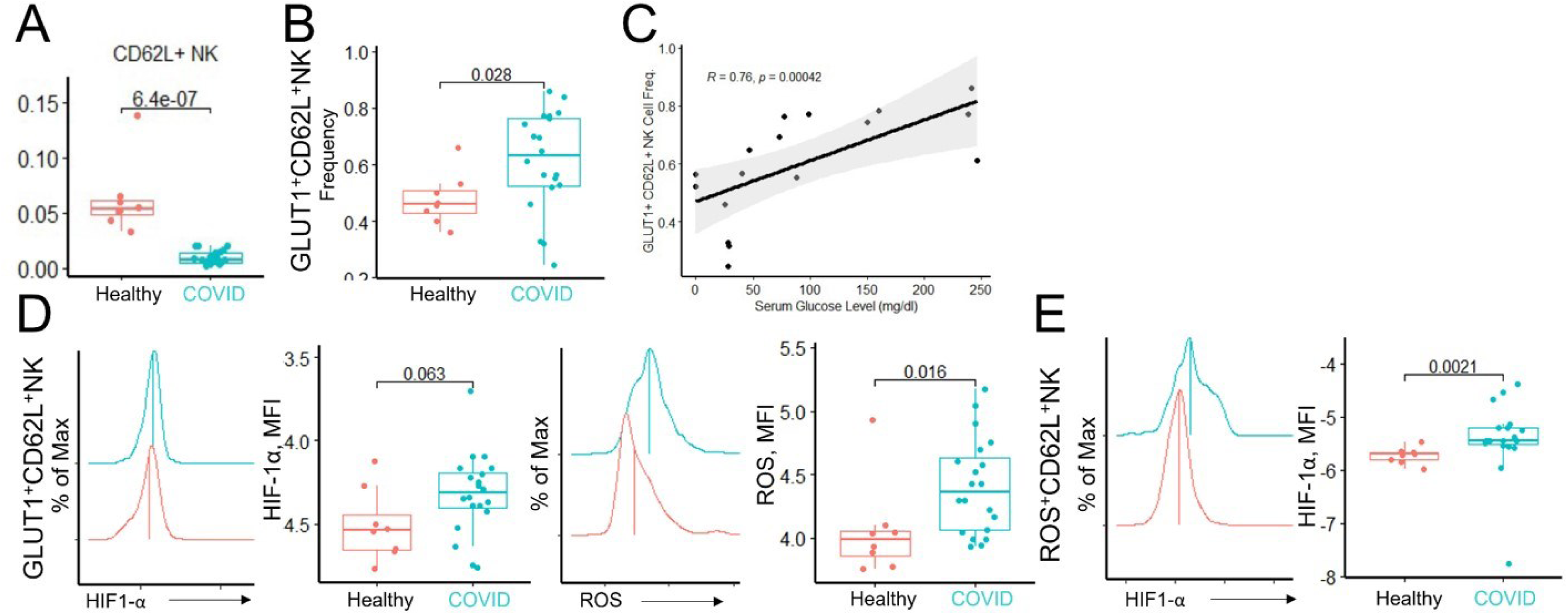
Aberrant metabolism of memory NK cells in COVID-19 patients. Impaired metabolic phenotype and differentiation of memory-like CD62L^+^ NK cells in circulation during COVID-19. **A.** Box-plot of cell-type proportion of CD62L^+^ NK cells for each disease state, each dot represents individual sample, 2-sided Wilcoxon Mann Whitney test was performed to indicate statistical significance; **B.** Box-plot of cell-type proportion of GLUT1^+^ CD62L^+^ NK cells for each disease state; **C.** Scatterplot demonstrating correlation between serum glucose level and frequency of GLUT1^+^ CD62L^+^ cells in COVID-19 patients, linear regression line with error bars displayed along with spearman correlation statistics; **D.** Density plot displaying distribution of HIF- 1a expression in GLUT1+ CD62L+ NK cells, adjacent **to** boxplot displaying MFI values of HIF-1a expression in GLUT1+ CD62L+ NK cells for both Healthy and COVID-19 patients; **E.** Density plot displaying distribution of ROS expression in GLUT1+ CD62L+ NK cells, adjacent to boxplot displaying MFI values of ROS expression in GLUT1+ CD62L+ NK cells for both Healthy and COVID-19 patients; **F.** Density plot displaying distribution of HIF-1α expression in ROS+ CD62L+ NK cells, adjacent to boxplot displaying MFI values of HIF-1a expression in ROS+ CD62L+ NK cells for both Healthy and COVID-19 patients;

### Metabolic dysregulation impairs immune surveillance and increases proinflammatory response in lung epithelial cells during SARS-CoV-2 infection

Epithelial cells (ECs) secrete cytokines and help mediate antigen presentation in order to modulate immune cells function during viral infection^33^. Differential expression analysis revealed overexpression of key immune signaling pathways in COVID-19 ECs (**Figs. 6A**). Network analysis demonstrates a connection of COVID-19 infection with the downregulation of the transcriptional factors, *ZKSCAN1* and *CSNK2B,* and upregulation of *KLF6, NEAT1,* and *JUND* (**Figs. 6B**). Induction of a pro-inflammatory cascade including type 1 IFN, toll-like receptor, NF-kB, and chemokine signaling and PI3K/AKT pathway was observed in COVID-19 ECs (**Figs. 6C,G**). Glucose metabolism mediates type I IFN secretion through enhancing transcriptional expression and epigenetic acetylation^34^. Indeed, we found a positive correlation between module scores for glycolysis and type 1 IFN signaling, as well as for glycolysis and NF-kB signaling (**Figs. 6D, E**). Chronic presentation of viral antigens to CD8 T cells by ECs may cause cellular dysfunction^35^. We observed that genes encoding HLA class 1 (*HLA-E, PSMA-6, TAP1, IFI30)* were enriched in COVID-19 ECs (**Fig. 6A**). GSEA analysis further confirmed the upregulation of HLA class 1 antigen presentation in bulk ECs. In contrast, downregulation of genes encoding HLA class 2 (*HLA-DRA, HLA-DPA1, HLA-DMA, DYNLL1)* was found in COVID-19 ECs (**Fig. 6A**) which was further confirmed by GSEA analysis (**Fig. 6F**). Glycolysis was reported to repress functional response of antigen presenting cells during infection^36^. We indeed observed a negative correlation of glycolysis and genes encoding HLA class 2 machinery (**Fig. 6H**). These results revealed potential links between dysregulated EC metabolism with cytokine release syndrome and adaptive immune dysfunction in COVID-19.

**Fig. 6.**
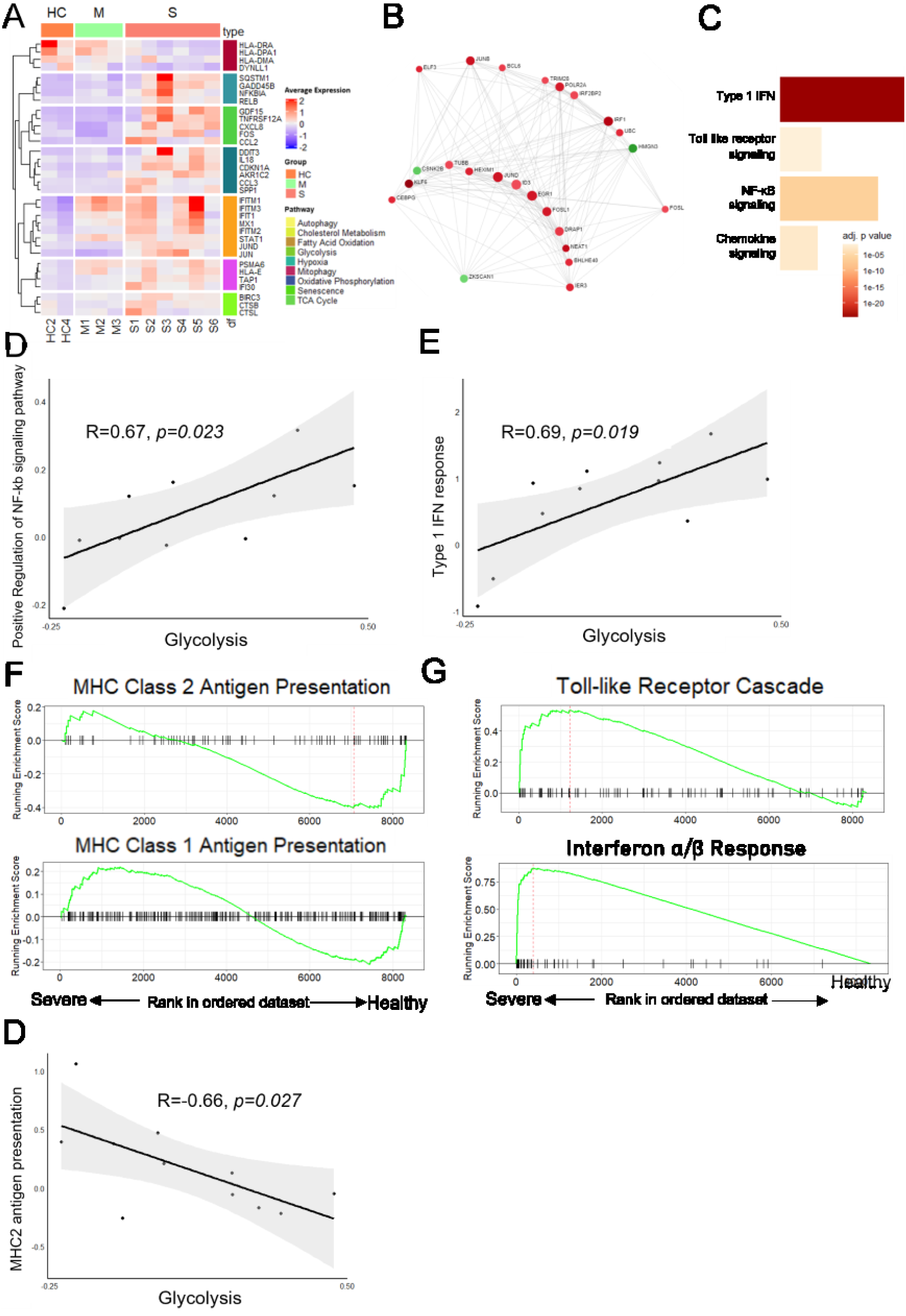
Metabolic disorder impairs immune surveillance and inflammatory response of ECs. The impact of metabolic reprogramming on EC immunological response was investigated. **A**. Heatmap displaying expression of key differentially expressed genes regulating immune signaling; **B**. Bar plot showing GSEA results of key statistically significant immune signaling pathways, x axis displays number of enriched genes per pathway, bars are colored by adjusted p. value; **C**. GSEA enrichment plots for “HLA Class 2 Antigen Presentation”, “HLA Class 1 Antigen Presentation”, “Toll-like Receptor Cascade”, and “Interferon A/B Response” pathways comparing severe vs healthy control patients for pseudostratified ciliated epithelial subset; **D**. Linear regression and pearson correlation analysis between module scores for glycolysis and HLA Class 2 Signaling; **E.** Linear regression and pearson correlation analysis between module scores for glycolysis and Type 1 interferon response; **F.** Linear regression and pearson correlation analysis between module scores for glycolysis and NF-kB signaling for all ECs; **G**. Network based display of transcription factor-gene interactions of differentially expressed genes between severe and healthy patients;

BALF ECs were next identified and subsetted for downstream analysis(**Fig. 7A**). Differential expression analysis revealed key differences in the expression of transcripts governing key metabolic pathways (**Fig. 7B**). Additionally, UMAP performed solely on differentially expressed metabolic genes revealed distinct clustering of bulk epithelial cells along disease severity (**Fig. 7C**). Pearson correlation analysis performed on ECs revealed a strong positive correlation between *HIF1A* and key glycolytic transcripts, suggesting a hypoxia-induced glycolytic metabolic reprogramming (**Fig. 7D**). ECs were then divided into pseudostratified ciliated and nonciliated subtypes based on the expression of canonical genes associated with cilia production (*CFAP126*, and *DNAAF*) (**Figs. 7E**). The ratio of pseudostratified ciliated ECs to nonciliated epithelial cells was inversely correlated with COVID-19 disease severity (**Fig. 7F**). This finding suggested that SARS-CoV-2 infection produced direct injury to the ciliated EC compartment. Overexpression of glycolytic transcripts *(ENO1, ADH1A3, GAPDH, ALDOA, PCK2)* was noted in both ciliated and nonciliated EC subsets from COVID-19 infected patients (**Fig. 7G**). These results were validated by GSEA analysis, which demonstrated enrichment of glycolysis genes (**Fig. 7G**). We also observed decreased expression of FAO regulating genes to different extent in ciliated and nonciliated ECs from S-COVID-19 compared to HC (**Fig. 7G**). *HIF-1A* and anaerobic glycolysis gene expression was strongly correlated with reduced expression of the OXPHOS and TCA cycle genes in these EC subsets from S-COVID-19 (**Fig. 7H**). GSEA analysis demonstrated enrichment of glycolysis, as well as a large downregulation of OXPHOS and TCA cycle regulating genes in S-COVID-19 ciliated and nonciliated ECs (**Fig. 7H**). Collectively, these results suggested that oxygen deprived conditions in the COVID-19 lung mediates a metabolic switch from aerobic FAO and OXPHOS towards anaerobic glycolysis in ECs, which is strongly linked to mitochondrial dysfunction.

**Fig. 7.**
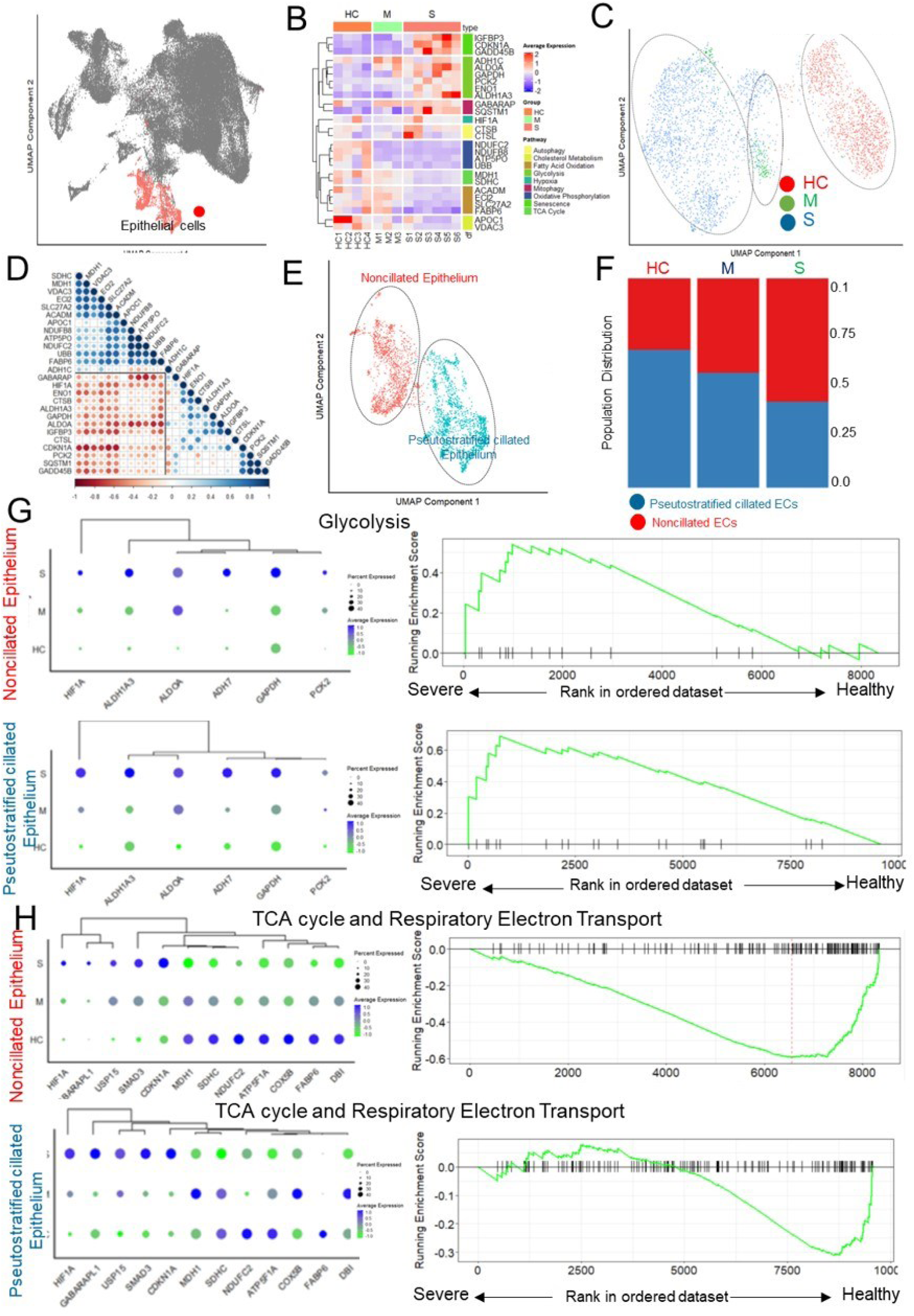
Metabolic dysregulation of BALF-derived ECs in COVID-19. Evaluation and assessment of metabolic phenotypes of epithelial subtypes shed in the BALF. **A** UMAP projection of 3531 ECs from Healthy Control (2), Moderate (3), and Severe (6) patients; **B.** UMAP projection of labelled epithelial cell subpopulations; **C.** Bar graph showing distribution of pseudostratified ciliated ECs and nonciliated ECs across conditions; **D.** Heatmap displaying expression of key differentially expressed metabolic genes; **E.** UMAP projection of ECs clustered solely on the expression of 42 differentially expressed metabolic genes; **F.** Correlation matrix showing spearman correlations between differentially expressed metabolic genes; **G.** Dot plots demonstrating expression and hierarchical clustering of select key glycolytic genes for ciliated ECs, adjacent is a GSEA enrichment plot for “Glycolysis” comparing ciliated ECs from severe vs healthy control patients; **H.** Dot plots demonstrating expression and hierarchical clustering of select key mitochondrial metabolism genes for ciliated ECs, adjacent is a GSEA enrichment plot for “TCA and Respiratory Electron Transport” comparing ciliated ECs from severe vs healthy control patients; **I.** Dot plots demonstrating expression and hierarchical clustering of select key glycolytic genes for nonciliated ECs, adjacent is a GSEA enrichment plot for “Glycolysis” comparing nonciliated ECs from severe vs healthy control patients; **J.** Dot plots demonstrating expression and hierarchical clustering of select key mitochondrial metabolism genes for nonciliated ECs, adjacent is a GSEA enrichment plot for “TCA and Respiratory Electron Transport”” comparing nonciliated ECs from severe vs healthy control patients

## Discussion

Metabolic syndrome and its accompanying effects are significant risk factors for COVID-19 lethality^5,37^. Understanding the role of cellular metabolism in COVID-19 pathogenesis has key implications in both COVID-19 prognosis and treatment. Here, we used single-cell omics techniques to construct a comprehensive metabolic landscape of immune cells involved in the SARS-CoV-2 anti-viral response. Evaluating cells from both the BALF and the blood of hospitalized COVID-19 patients revealed that lymphocyte populations undergo a global metabolic reprogramming towards anaerobic processes, resulting in compromised memory cell differentiation and effector function. We were also able to identify and define highly resolved states of dysregulated cellular metabolism in key immune cell subsets that could potentially indicate COVID-19 severity.

Despite ARDS being the main contributor to mortality and severe complications during COVID- 19 infection^38^, prior attempts to elucidate the role of metabolism in lymphopenia and immune cell exhaustion during severe COVID-19 are limited to solely analysis of patient-derived PBMCs^8,39,40^. Cossarizza *et al.* used flow cytometry and metabolic flux assays to characterize the immunometabolic phenotype of isolated PBMCs from COVID-19 patients^40^. Although an increased abundance of exhausted PD1^+^ lymphocytes in severe patients was found, this was not accompanied by any difference in extracellular acidification (ECAR) and oxygen consumption rates (OXPHOS)^40^. Additionally, using flow cytometry and single cell RNA-sequencing, the Powell group discovered a novel population of VDAC1^+^ exhausted T cells in PBMCs from severe COVID- 19 patients ^41^. However, they did not detect any significant change in the expression of glycolysis-regulating genes in T cells ^41^, which is contradictory to our results. One potential explanation for this discrepancy is that the aforementioned study stimulated isolated PBMCs with αCD3/CD28 polyclonal activation or with SARS-CoV-2 specific peptide libraries. Prolonged maintenance and stimulation of T cells under normoxic conditions is not reflective of the hypoxic microenvironment conditions present during severe COVID-19 infection, which may result in the attenuation or nulling of any potential metabolic differences present in COVID-19. Another potential explanation arises from the fact that cellular metabolism in these studies was assessed for entire T cell populations. Given that T cells are highly heterogeneous with respect to their metabolism, metabolic profiles need to be assessed on specific T cell subpopulations instead of bulk T cells. In contrast, by using high-dimensional flow cytometry to investigate the single-cell metabolism of unstimulated PBMCs immediately after isolation, and assessing single cell transcriptomic data from the BALF, we aimed to capture the metabolic state of the cells from their original microenvironment. Our single-cell omics approach allowed us to detect distinct differences in metabolic phenotypes of specific, highly resolved lymphocyte populations. We found that in response hypoxic conditions, there is a metabolic reprogramming of CD8 and NK cell subsets traditionally reliant on OXPHOS and FAO toward anaerobic glucose metabolism, which along with mitochondrial dysfunction, triggers cellular exhaustion and disrupts memory differentiation, resulting in a compromised anti-viral response.

In this study, we uncover a clear link between cellular metabolism, severe COVID-19, and disrupted memory cell development. During infection, hypoxia resulting from lung epithelial cell damage triggers a global metabolic reprogramming of CD8 and NK cell subsets from OXPHOS and FAO towards anaerobic glucose metabolism. Upon activation, CD8 cells typically exhibit a Warburg like metabolic adaptation to rely on aerobic glycolysis; thus, a metabolic switch from aerobic to HIF-1a mediated anaerobic glycolysis will not impair initial T cell activation into effector subsets, which is consistent with reports demonstrating hyperactivation of CD8 T cells^42^. However, because of prolonged anaerobic glycolysis, the lung microenvironment becomes increasingly hostile, resulting in nutrient depleted, hyperlactatemic, and hypoxic conditions. Herein, we show that this dysregulated metabolism is heavily tied with impaired memory lymphocyte formation. Although memory lymphocytes are traditionally associated with a heavy reliance on OXPHOS and fatty acid oxidation, we have detected multiple clusters of cellular state associated with high glucose uptake, ROS production, hypoxia mediated transcriptional response, and cellular exhaustion in CD8 and NK memory cell populations that were specifically enriched in hospitalized COVID-19 patients. These populations were negatively associated with memory lymphocyte frequency, suggesting that this metabolic switch may halt the transition of activated lymphocytes into memory cells. Pseudotime analysis and trajectory interference with transcriptomic data further demonstrate differentiation of tissue-resident memory cells that was tied to altered lymphocyte metabolism. In addition to anaerobic glycolysis, upregulation of glutaminolysis and mitophagy was also seen in CD8 memory cells in order to sustain bioenergetic demands. Clustering solely on metabolic phenotype revealed clear distinctions between healthy, moderate, and severe patients in CD8 memory cells, suggesting the potential use of metabolic markers in predicting memory cell response.

Interestingly, we found that SARA-CoV-2 derived EC damage creates oxygen-deprived conditions in the lungs that not only induce metabolic reprogramming of various immune cell subsets, but also themselves. We found that during COVID-19 infection, differential metabolism drives lung ECs towards senescence and towards acquiring a significant SASP phenotype, leading to secretion of proinflammatory cytokines, reduced HLA class 2 mediated immunosurveillance, and increased HLA class 1 machinery. Chronic stimulation of exhausted lymphocytes, that demonstrate attenuated effector function and cytokine secretion in nutrient-depleted microenvironments, by antigen presenting cells via HLA class 1 leads to significantly increased cellular exhaustion, which further impairs the capacity of cells to differentiate into memory phenotypes. Our results therefore show that the immunometabolic rewiring of ECs in the BALF can be a potential mechanism for organ-specific lymphocyte exhaustion and memory cell dysfunction. Further, this observation thus highlights that unconventional antigen presentation on non-hematopoietic ECs via HLA class 1, in addition to conventional antigen presentation by professional APCs (monocyte, DC, and macrophage), can be considered as a potential target for therapeutic development.

Reports have shown that unconventional T cells such as NKTs are attenuated in COVID-19^43^. However, the mechanisms underlying these observations are unknown. Herein, we provided evidence suggesting that ECs induce NKT exhaustion and dysfunction through prolonged antigen stimulation resulting from a hypoxia-mediated metabolic adaptation. The traditional view that NK cells are short-lived innate lymphocytes is being challenged by new data demonstrating that NK cells can develop long lasting, antigen- specific memory in response to viral infection^44^. Recently, CD62L^high^ NK were identified as a subset possessing multiple characteristics of memory cells, demonstrating rapid responsiveness towards viral restimulation^32^. In the current study, we discovered that metabolic disorders cause CD62L^+^ NK lymphocytopenia via impairment of memory formation. Because the abundance of GLUT-1^+^CD62L^+^NK cells can predict the COVID- 19 severity, further studies about the role of CD62L^high^ NK cells in COVID-19 pathogenesis are crucial.

Targeting T cell glycolysis during COVID-19 infection might be a promising approach for rescuing T cell fate and function. Ideally, we suggest that attempts to target T cell glycolysis in COVID-19 should take place after clonal expansion and formation of initial antigen specific T cells, but before initiation of memory cell formation. It is possible that the efficacy of dexamethasone treatment in COVID-19 patients on mechanical ventilation and supplemental oxygen ^45^ may be due to steroid-mediated inhibition of glycolysis^46^. Thus, dexamethasone administration may be able to rescue T cell dysfunction, improve memory cell formation, and reduce CRS via inhibition of glycolysis^47^. A recent stage-2 clinical trial reported that the use of 2-DG, a glucose analogue used to inhibit glycolytic flux, as a therapeutic treatment for COVID-19 was highly successful in improving patient outcomes, furthering glycolysis as a potential target to restore T cell fate^48^. In support of the importance of time-dependent treatment for COVID-19, type 1 IFN, which was reported to induce the metabolic reprogramming from glycolysis into OXPHOS and FAO in immune cells^49^, was only effective as a treatment option in COVID-19 when administrated early after infection^50^. In contrast, delayed type 1 IFN treatment resulted in worsening of COVID-19 severity due to the pro-inflammatory induction capacity of this cytokine ^50^. Further, our results implicate mitophagy as a potential target for therapeutic intervention. In response to a nutrient-depleted and hypoxic microenvironment, effector CD8 T cells may upregulate mitophagy as an alternative survival mechanism. However, prolonged upregulation may induce lymphocyte exhaustion and mitochondrial dysfunction. Accordingly, ablating mitophagy in CTLs can potentially redirect T cells towards memory cell differentiation and rescue them from exhaustion^51^. Furthermore, unlike glycolysis, mitophagy is not critical for T cell activation and effector cell differentiation^51^. Thus, mitophagy- targeting approaches can potentially be used immediately after COVID-19 infection. Targeting mitophagy with a specific mitophagy inhibitor such as Mdivi-1 should be explored as a treatment regimen for COVID-19.

Supplemental oxygen as opposed to ventilation was shown to improve the outcome of patients with severe COVID-19^52^. Our current study suggests that hypoxia is a key regulator of immunometabolic dysfunction during severe SARS-CoV-2 infection. Hence, efforts to maintain blood oxygen saturation early in the course of infection are vital for patient recovery and improvement. Immunological outcomes post- initial infection such as memory cell formation dictate the severity of response upon re-exposure to the virus^35^. In this regard, metabolism can also influence the immune response after recovery from SARS-CoV-2. Because both effector function and memory differentiation are severely impaired in lymphocytes, T cell immunity may be compromised even in recovered patients. Given the cumulatively rising COVID-19 reinfection rate^53^, future studies investigating how metabolism affects the humoral response, including activation, antibody secretion, and long-term plasma cell differentiation of memory B, are imperative. We could not detect any population of T follicular helper cells in the BALF from 13 severe patients, suggesting that metabolic dysregulations also impaired germinal center formation in the lungs of COVID-19 patients. In support of our current observation, hypoxia and nutrient deprivation are known to suppress the generation of germinal center B cells and follicular helper cells after viral infection^54^. Additionally, defective Tfh and germinal B cell formation in spleen and lymph node, along with SARS-CoV-2-specific B cell enrichment in blood of severe COVID-19 patients, were recently reported ^55^. Overactivation of extrafollicular B cells in COVID- 19 is also indicative of germinal center impairment^56^. These findings suggest that patients with metabolic comorbidities or ARDS may suffer from a limited durability of antibody responses during COVID-19 infection. Furthermore, knowledge of how cellular metabolism regulates memory cell differentiation may help predict the reaction of patients with preexisting metabolic comorbidities to vaccination. Although COVID-19 vaccines that have been approved for emergency use and their short term effectiveness has been validated, their effectiveness in inducing long term immunity has yet to be established^57,58^. Furthermore, there has yet to be an attempt to understand the longevity of convalescence-induced protective immune responses in COVID-19 patients with metabolic disorders.

Despite claims of biomarkers to predict COVID-19 severity^59^, no specific markers for COVID-19 patients with metabolic co-morbidities have been yet discovered. In the current study, using high dimensional analyses, we provide a number of lowly abundant cell populations in the blood of severe COVID-19 patients that can potentially predict disease severity including GLUT1^+^ mitochondrially exhausted CTL, CD8CM, NKT and NK cells. Noticeably, a clear correlation between the serum glucose level, recently identified as risk factor COVID-19 severity in patients with pneumonia^60^, and GLUT-1^high^ CD62L^+^ NK cells was observed, suggesting that the use of metabolic biomarkers in combination can be strong prognostic indicator for COVID-19 disease severity.

Overall, our current study sheds important new light on the molecular and cellular mechanisms by which immune cell metabolism regulates COVID-19 pathobiology. Shortcomings of our study include a limited sample size for analysis of BALF transcriptomic data, along with a lack of proteomic data for our study of lung ECs. Additionally, our assessment of cellular metabolism is limited to analysis surface and intracellular metabolic marker expression, coupled with transcriptomic data for key metabolic pathways. Future studies should aim to validate our results using approaches that shed more light into functional metabolism such as Seahorse Metabolic Flux profiling or metabolomics. Given the large diversity in the range of responses towards SARS-CoV-2 infection, stratification for factors such as gender, BMI, preexisting conditions, and prior treatments would help to further validate our analysis. Nonetheless, this study provides crucial information about the pivotal relationship between cellular metabolism and the memory lymphocyte response during severe COVID-19. Our results shed light on novel biomarkers, therapeutic targets, and strategies for the COVID-19 therapy. We also provide a novel data analysis pipeline for understanding single cell metabolism in an organ-specific manner. We are confident that this comprehensive single-cell transcriptomic and proteomic portrait of immune cell metabolism will advance COVID-19 research and help to devise novel approaches for mitigating the COVID-19 pandemic.

## Methodology

### Sample Acquisition

Blood from healthy donors were ordered from Research Blood Company. Blood samples from hospitalized COVID-19 patients were collected from the AdventHealth hospital under protocols IRB# 1668907 and #1590483 approved by AdventHealth IRB committee. Strict confidentiality was maintained for all patients according to HIPAA confidentiality requirements. COVID-19 was confirmed by PCR test at AdventHealth. Blood was used for human PBMC, plasma, and serum isolation.

### PBMC Isolation

PBMCs were isolated by density-gradient centrifugation using Ficoll. Briefly, blood specimens were centrifuged at 700G for 7 min at RT for serum collection. The pellets were resuspended in phosphate buffer saline (PBS). Cell suspension were carefully overlay on the top of 4 mL Ficoll in 15 mL conical tube, followed by centrifugation at 700G at RT for 25 min without break. PBMCs were collected from interphase between plasma and Ficoll layers. Cells were wash twice with PBS to remove Ficoll residue. All the procedures were approved at BSL2^+^ level by UCF Environmental Health and Safety.

### Antibody Staining and Flow Cytometry

About 5×10^5^ cells from each sample were used for flow cytometry staining. See Table S3 for antibody information. PBMCs were first stained with live/dead in PBS for 15 min, washed with FACS buffer, and stained with surface markers in FACS buffer at 4°C for 30min. Following incubation, PBMCs were washed (FACS buffer, 200 µL), stained with secondary antibody mix for another 15 min, and then washed again with FACS buffer. Samples were fixed and permeabilized with Fixation/Permeabilization buffer (15 min) and washed with FACS buffer. PBMCs were then stained with the intracellular antibody staining mix at 37°C for 45 min. Sample were washed once with Fixation/Permeabilization buffer before being resuspended in FACS buffer for flowcytometric analysis.

### High-dimensional Flow Cytometry Analysis

First, the flowCore package in R was used to read in compensated FCS files into the R environment^61^. Automatic gating functions in openCyto were next used to filter cells for doublets and debris^62^. Next, an arcsinh transformation with a cofactor of 5 was applied for data normalization. Data from all of the samples were then merged into one Catalyst object, upon which downstream analyses were performed^63^. CytoNorm was next applied to correct for batch effect between samples by aligning peaks of bimodally distributed markers^64^. PCA was then run on bulk sample-aggregated data and the top 3 principal components were plotted. FlowSOM clustering was performed on only cell surface markers used for phenotypic identification with the number of expected populations set at 20^65^. Clusters were then annotated based upon canonical marker expression. For purposes of UMAP dimensionality reduction, clusters that could not be labelled were discarded before UMAP was performed. Differential abundance of cell-type proportions and differential expression of MFI values were next conducted. For each identified population of interest, FlowSOM was applied again on only the functional state markers with the number of expected populations set at 10. Unsupervised clusters were then annotated by canonical marker expression to define highly -resolved cell states for specific populations. Briefly, after quality control and compensation, batch effect was corrected in makers with bimodally distributed expression by the CytoNorm package in R^64^. UMAP and FlowSOM^65^ were used to identify unsupervised clusters (**Fig. 1H** and **Fig. S3A**) which were assigned to specific populations based on canonical marker expression (**Fig. 1I** and **Fig. S3B**).

### BALF Data Acquisition

Single cell RNA-seq data from the BALF of 6 severe COVID patients, 3 moderate patients, and 4 healthy donors were used for analysis ^66^. This study defined moderate and severe COVID-19 patients as those with pneumonia experiencing respiratory distress and hypoxia and with critical condition, requiring ICU care, and having been placed under mechanical ventilation, respectively. Prefiltered expression matrices with UMI counts were downloaded from the GEO Database with accession number GSE145926. Additionally, as suggested by the original study, data from an additional BALF sample derived from a healthy donor from a separate study was used as a reference ^67^. Prefiltered expression matrices with UMI counts were downloaded from the GEO Database with accession number GSE128033 and sample number GSM3660650.

### Data Quality Control and Preprocessing

Quality control and data preprocessing was done using Seurat ^68,69^. First, cells for which more than 10% of reads were mitochondrial transcripts were discarded. Next, we removed cells that had less than 1000 detected transcripts. Cells with less than 200 and greater than 6000 unique genes were also filtered. Filtered data from different 14 patient samples were integrated in Seurat. Individually, data from each sample was log 2 normalized and the top 2000 variable genes were identified using the “vst” method in Seurat. Data from each sample was next scaled and PCA was run with percentage of mitochondrial DNA and number of detected unique genes regressed out. Alignment and batch effect correction was done using reciprocal PCA and canonical correlation analysis (CCA) (in accordance to standard Seurat integrated analysis workflow) on the first 30 dimensions of the data. Next, a shared nearest neighbor graph was constructed and Louvain-based optimization was run to perform unsupervised clustering. UMAP was next run on the first 30 dimensions. Data was next log 2 normalized and scaled in the “RNA” assay for expression analysis, with percentage of mitochondrial DNA and number of detected unique genes regressed out. The top 2000 variable genes were determined by the “vst” method in Seurat. Expression of canonical markers were used to define cell populations. For each identified cell population, SCTransform was done on the “RNA” assay to improve normalization and aid in visualization purposes.

### T cell reintegration

T cells were subsetted and split according to samples. Data from healthy control 1 and severe 1 were excluded from analysis due to low T cell count. To further correct for batch effect, T cells were then reintegrated using canonical correlation analysis in Seurat run on the first 30 dimensions. SCTransform was next implemented on the “RNA” assay and stored in a new “SCT” assay to better normalize counts across samples for visualization purposes with percentage of mitochondrial DNA regressed out. Standard log 2 normalization and scaling was then performed on the “RNA” assay. Subpopulations of T-cells were next identified based upon canonical marker expression.

### Trajectory Inference and Pseudo-temporal Ordering

Monocle 3 was used to construct a trajectory upon UMAP embeddings and order cells in pseudotime ^70^. Analysis was done on both CD8 and CD4 T cells. Seurat wrapper function “asMonocle” was used to create Monocle CellDataSet object from an existing Seurat object. “learn_graph” function was used to construct trajectory mappings onto transferred UMAP embeddings. “order_cells” was used to estimate and order cells in pseudotime. All samples for CD8 and CD4 populations were ordered together and were split by disease state after ordering for differential comparison of pseudotime.

### Metabolic Phenotype based Clustering

To investigate whether metabolic phenotypes of certain cell populations alone could be used alone as predictive indicators of disease severity, dimensionality reduction at both a single cell and sample-wide resolution was done only on key identified differentially expressed metabolic genes to see if cells/samples would cluster according to disease severity. For sample-wide analysis, principal component analysis was conducted and the first three principal components were visualized. For analysis at single cell resolution, UMAP was done and the first two components were visualized.

### Network Analysis

For construction of gene pathway enrichment network, networkanalyst.ca was used ^71^. All statistically significant genes were inputted along with log fold change values to construct enrichment network. Transcription factor – gene interaction network was also constructed using networkanalyst.ca ^71^. Statistically significant genes along with log fold changes vales were inputted. The “degree” filter was first set to 100 and then the “betweenness” filter was set to 170.

### Downstream Analysis

For heatmap visualizations, scaled SCTransformed values were used and the Complexheatmap package was used to generate visualization ^71^. Hierarchical clustering and dendrogram generation were done using default settings of the package. Outliers with extremely high scaled expression values (>> 2) were set to a maximum value of 2 to not distort the rest of the Fig. . For dotplot visualizations, first a euclidean distance matrix was generated for which hierarchical clustering was then applied. Ggtree was next used for dendrogram construction ^72^. ReactomePA package was used for functional GSEA ^73^. All unique detected genes in the cell subset were sorted by log fold change values to create ranked list that was inputted for GSEA analysis. enrichR was used to determine over and under expressed pathways from differential expression analysis (Kuleshov) ^74^. Corrplot package was used for generation of correlation matrices. Volcano plots were constructed using EnhancedVolcano. Other graphical visualizations were created using ggplot2, ggpubr or plotly. All further downstream analysis was done in base R.

### Statistical Analysis

Differential expression analysis of transcript abundance was assessed using Seurat’s implementation of the nonparametric Wilcoxon rank-sum test. Genes were generally defined as statistically significant by Bonferroni adjusted p. values less than 0.05 and log-fold change greater than 0.25. For NKT cells, non-adjusted p. value was used to define differentially expressed genes due to very small sample size.

For comparison of cell-type proportions, nonparametric Wilcoxon rank-sum test was performed, and non-adjusted p. values were used to indicate significance. For comparison of median fluorescent intensity (MFI), nonparametric Wilcoxon rank-sum test was also used to evaluate significance. Additionally, Pearson correlation coefficient was used to indicate strength of measured correlations. Student’s *t test* was used to evaluate significance of measured correlation.

## Graphical Summary

Attack of lung ECs by SARS-CoV-2 causes pulmonary damage leading to generalized hypoxia, which in turn increases anaerobic glycolytic metabolism. Concomitant changes of metabolic processes with increased MHC class I expression in lung ECs (ECs) drives immune cell exhaustion in CD8 and NK cells. An accompanying effect is interference with immune memory cell formation, which increases the vulnerability of the host to reinfection by SARS-CoV-2. Therefore, protecting immune cell function in the face of SARS-CoV-2 infection by targeting dysregulated immunometabolism is a promising approach to treat COVID-19 patients and improve clinical outcome.

## Code Availability

The source code used to reproduce our analysis can be accessed upon request from the corresponding author.

## Ethics Declaration

This study was approved by the IRB committee of AdventHealth hospital and UCF under protocols #1668907 and #1590483.

## Author contribution

H.N. conceived the research project. S.G. and H.N performed data analysis, drafted manuscript and wrote the paper. HN, SG, STG, DL, NTL contributed to revise the manuscript. HN, SG, AA, KE, DL, TH, SA, and AM contributed to the experimental process.

## Acknowledgements

The current study is conducted with the support from the University of Central Florida start-up funding and College of Medicine, University of Central Florida COVID-19 seed funding support to H.N. We would like to thank Drs. Alessandro Sette and Daniela Weiskopf at La Jolla Institute for Immunology for sharing COVID-19 peptide mix and Dr. Justine Tigno-Aranjuez for sharing flow antibody. Reagents and protocol for detection of COVID-19 antigen by ELISA were obtained through BEI Resources, NIAID, NIH.

## Abbreviations

COVID-19: coronavirus disease 2019
SARS-CoV-2: severe acute respiratory system coronavirus 2
BALF: bronchoalveolar lavage fluid
PMBC: peripheral blood mononuclear cells
FAO: fatty acid oxidation
CRS: cytokine released syndrome
CTL: cytotoxic T lymphocyte
SASP: senescence- associated secretory phenotype
EC: epithelial cells
Trm: T resident memory
ARDS: acute respiratory disease syndrome
p38 MAPK: p38 mitogen-activated protein kinase
LDHA: lactate dehydrogenase
PCA: principal component analysis
UMAP: universal manifold approximation and projection
CCA: canonical correlation analysis
ICU: intensive care unit

---

**Fig. S1.**
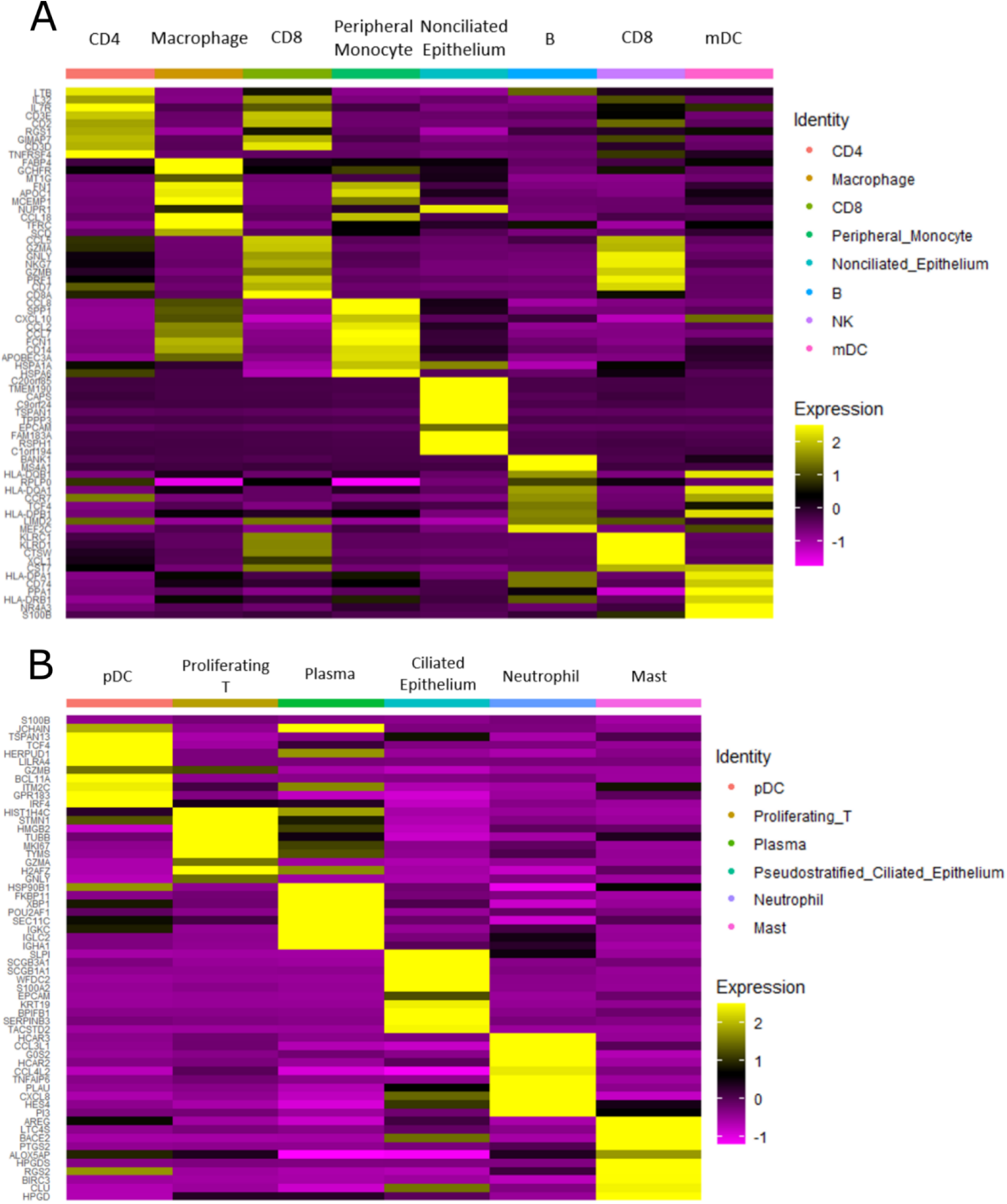
Heatmap displaying canonical gene expression of genes used to annotate unsupervised clusters for all 66,452 cells after sample integration. FindAllMarkers function in Seurat was used to identify the top genes specific to each annotated cell population A. CD4, Macrophage, Peripheral Monocyte, Nonciliated Epithelium, B, CD8, mDC B. pDC, Proliferating T, Plasma, Ciliated Epithelium, Neutrophil, and Mast). A heatmap displaying the average expression of the genes for each cell subset was generated using the DoHeatmap function in Seurat.

**Fig. S2.**
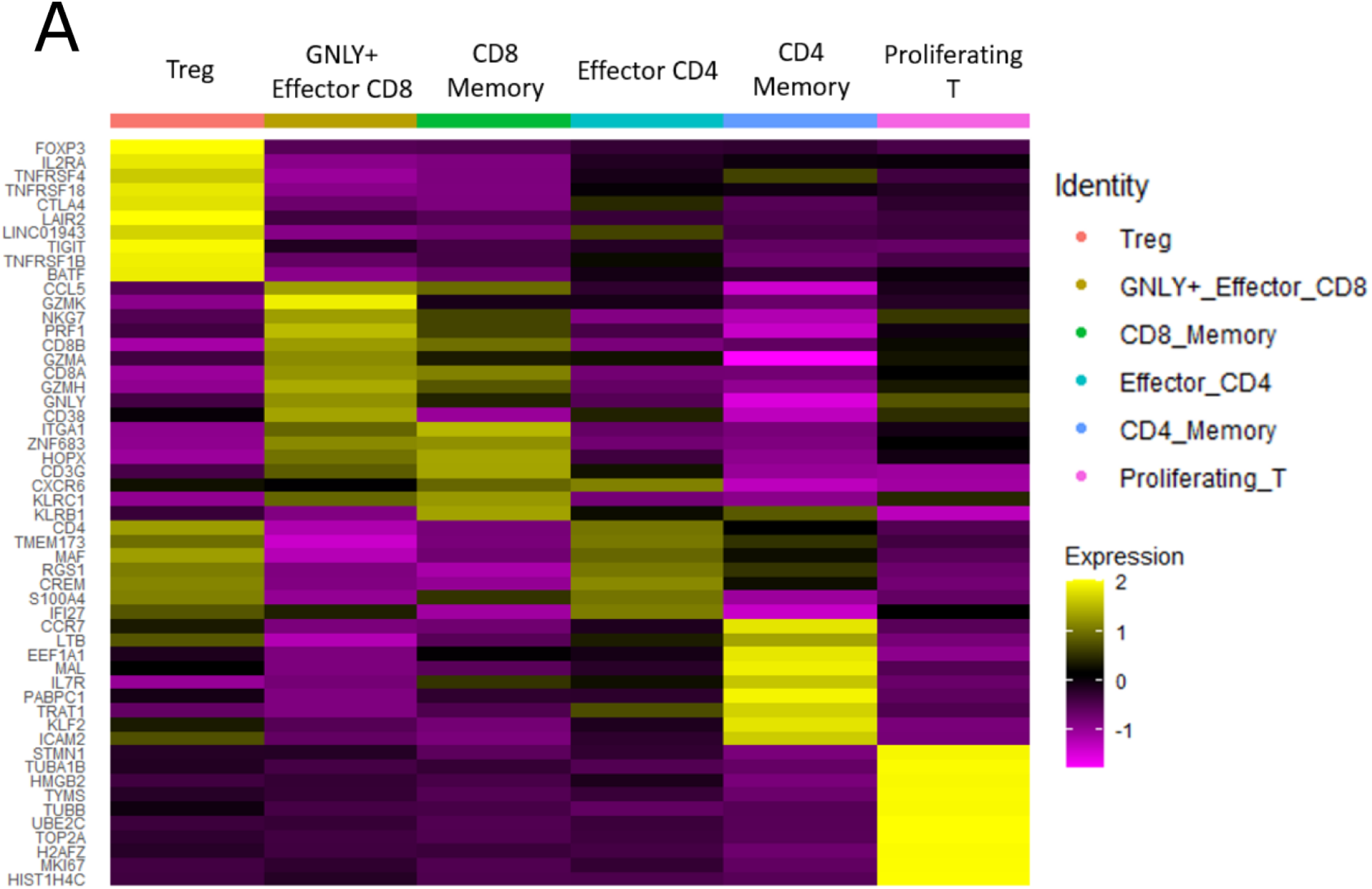
Heatmap displaying canonical gene expression of genes used to annotate unsupervised clusters after T cell reintegration. FindAllMarkers function in Seurat was used to identify the top genes specific to each annotated T cell population. The average expression of the genes for each T-cell subset was generated using the DoHeatmap function in Seurat.

**Fig. S3.**
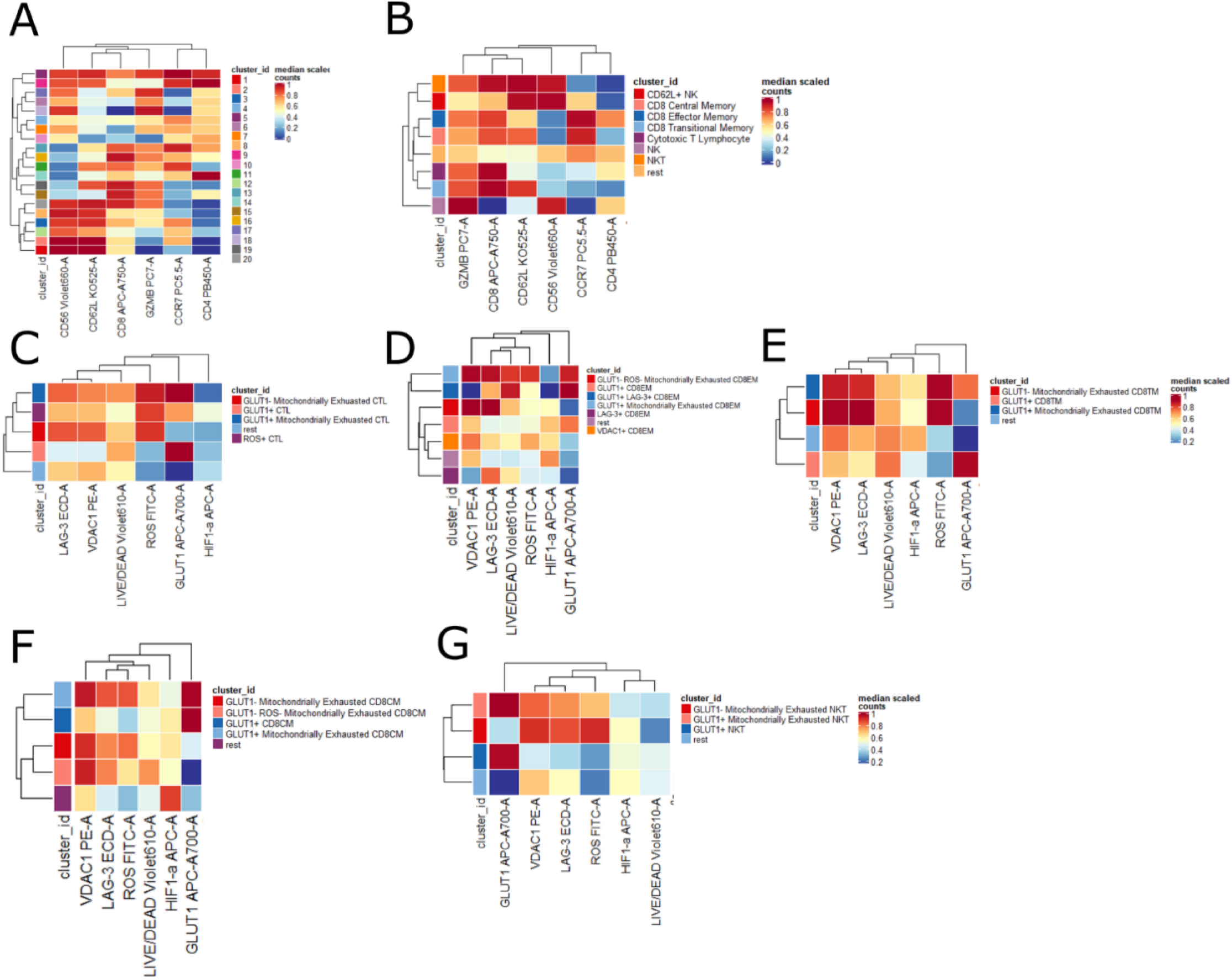
Heatmap displaying canonical marker expression of phenotypic markers. For unsupervised clusters generated through flowSOM (**A**); For annotated populations of PBMCs (**B**); CTLs in PBMCs (**C**); CD8EMs in PBMCs (**D**); CD8TMs in PBMCs (**E**); CD8CMs in PBMCs (**F**); NKT cells in PBMCs (**G**).

**Fig. S4.**
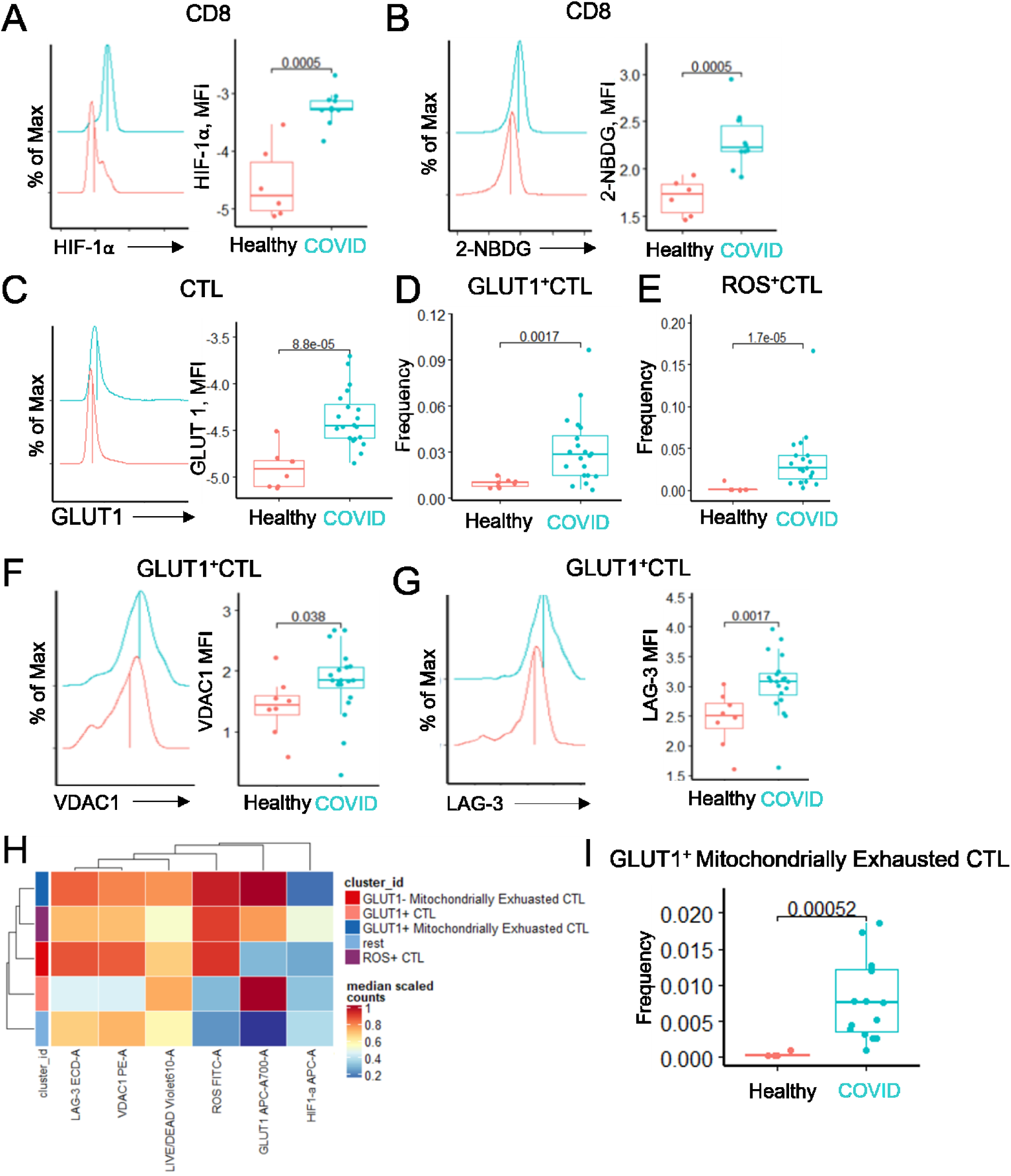
Metabolic profiles of circulating CTLs in COVID-19. CTLs identified by unsupervised clustering from patient PBMCs were evaluated to assess differential metabolic profiles. **A.** Density plot displaying distribution of HIF-1a expression in bulk CD8 cells, adjacent to boxplot displaying MFI values of HIF-1a expression in bulk CD8 cells for both Healthy and COVID-19 patients, each dot represents individual sample, 2-sided Wilcoxon Mann Whitney test was performed to indicate statistical significance; **B.** Density plot displaying distribution of 2-NBDG expression in bulk CD8 cells, adjacent to boxplot displaying MFI values of HIF-1A expression in bulk CD8 cells for both Healthy and COVID-19 patients. **C.** Density plot displaying distribution of GLUT-1 expression in CTLs, adjacent to boxplot displaying MFI values of GLUT-1 expression in CTLs both Healthy and COVID-19 patients; **D.** Box-plot of cell-type proportion of GLUT1+ CTLs for each disease state; **E.** Box-plot of cell-type proportion of ROS+ CTL for each disease state; **F.** Density plot displaying distribution of VDAC1 expression in GLUT1+ CTLs, adjacent **to** boxplot displaying MFI values of VDAC1 expression in GLUT1+ CTLs for both Healthy and COVID-19 patients. **G.** Density plot displaying distribution of LAG-3 expression in GLUT1+ CTLs, adjacent to boxplot displaying MFI values of LAG-3 expression in GLUT1+ CTLs for both Healthy and COVID-19 patients. **H**. Heatmap displaying canonical expression of labelled populations from the second round of unsupervised clustering; **I.** Box-plot of cell-type proportion of GLUT1+ Mitochondrially Exhausted CTLs for each disease state

**Fig. S5.**
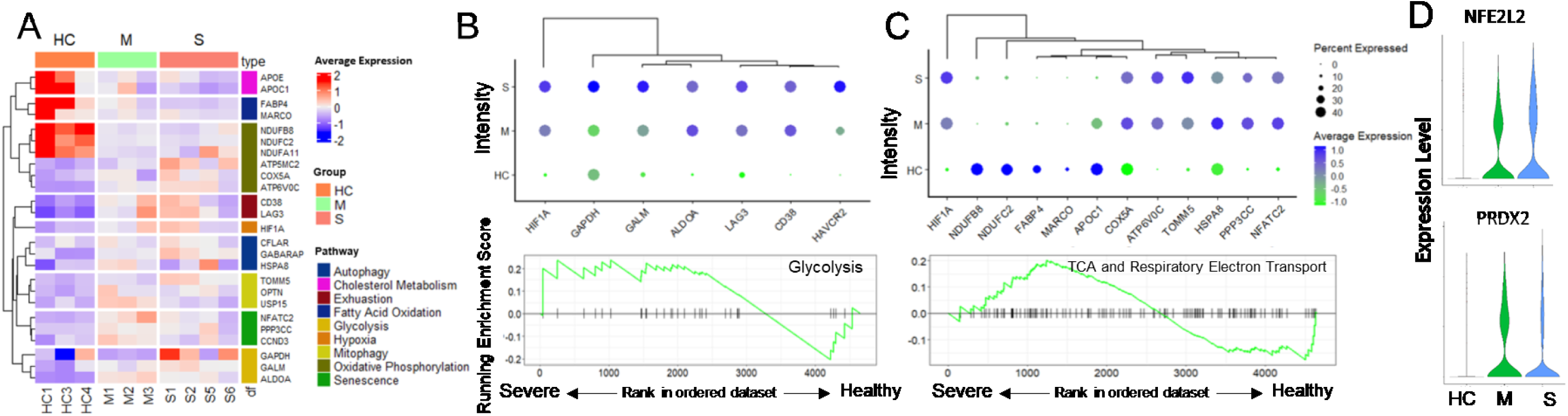
Differential metabolic phenotype of BALF derived CTLs in severe COVID-19. **A**. Heatmap displaying expression of key differentially expressed metabolic genes for CTLs; **B**. GSEA enrichment plots for “Glycolysis” comparing severe vs healthy control patients, adjacent is a dotplot demonstrating expression and hierarchical clustering of select key glycolytic genes; **C.** GSEA enrichment plots for “TCA and Respiratory Electron Transport” pathways comparing severe vs healthy control patients, adjacent is a dotplot demonstrating expression and hierarchical clustering of select key genes involved in mitochondrial metabolism; **D.** Violin plot demonstrating expression of NFE2L2 and PRDX2 across disease states.

**Fig. S6.**
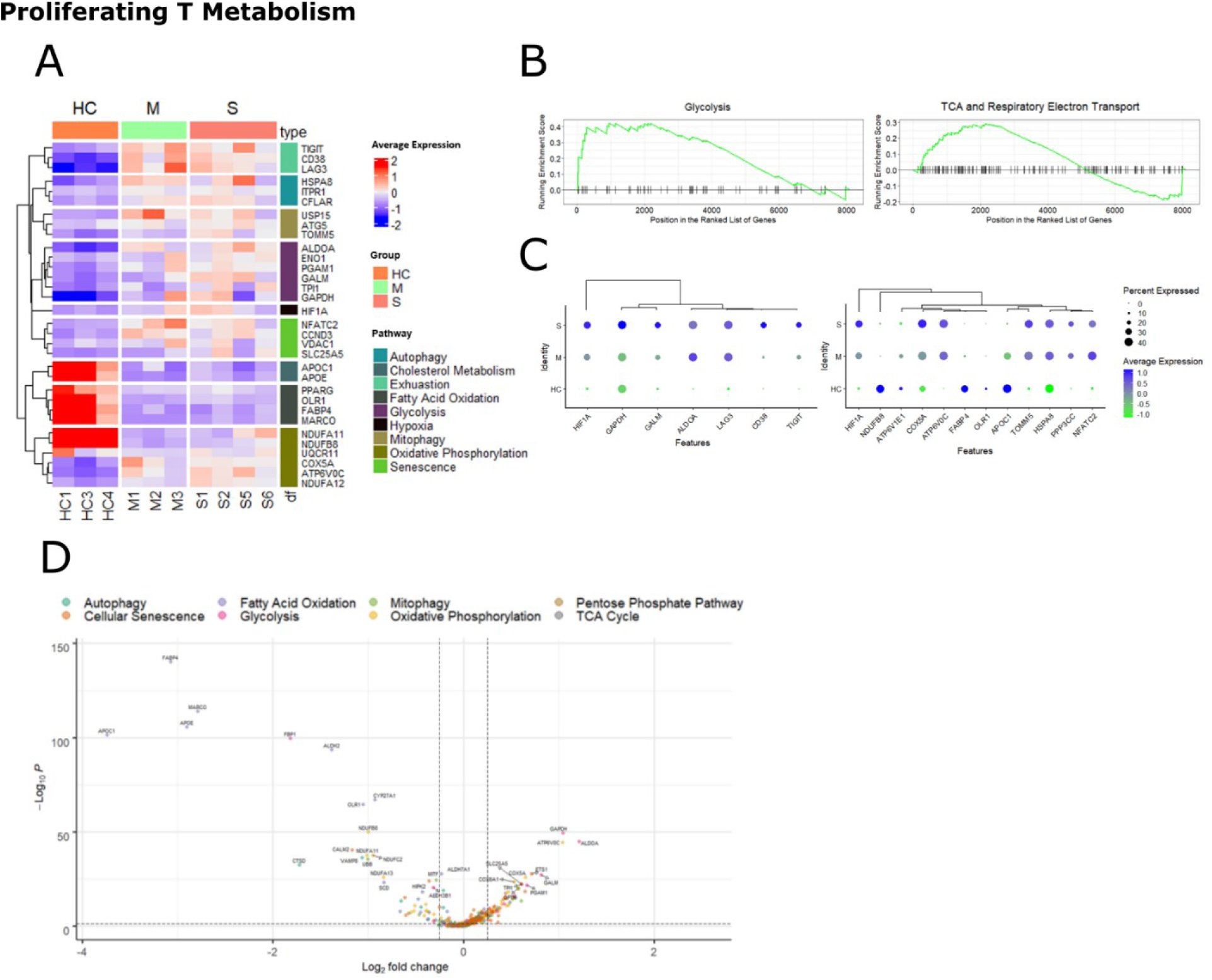
**A**. Heatmap displaying expression of key differentially expressed metabolic genes; **B**. GSEA enrichment plots for “Glycolysis” and “TCA and Respiratory Electron Transport” pathways comparing severe vs healthy control patients; **C**. Dot plots demonstrating expression and hierarchical clustering of select key metabolic genes; **D**. Volcano plot of differentially expressed genes between severe and healthy COVID-19 patients, x axis shows log2 fold change, y axis shows adj. p value

**Fig. S7.**
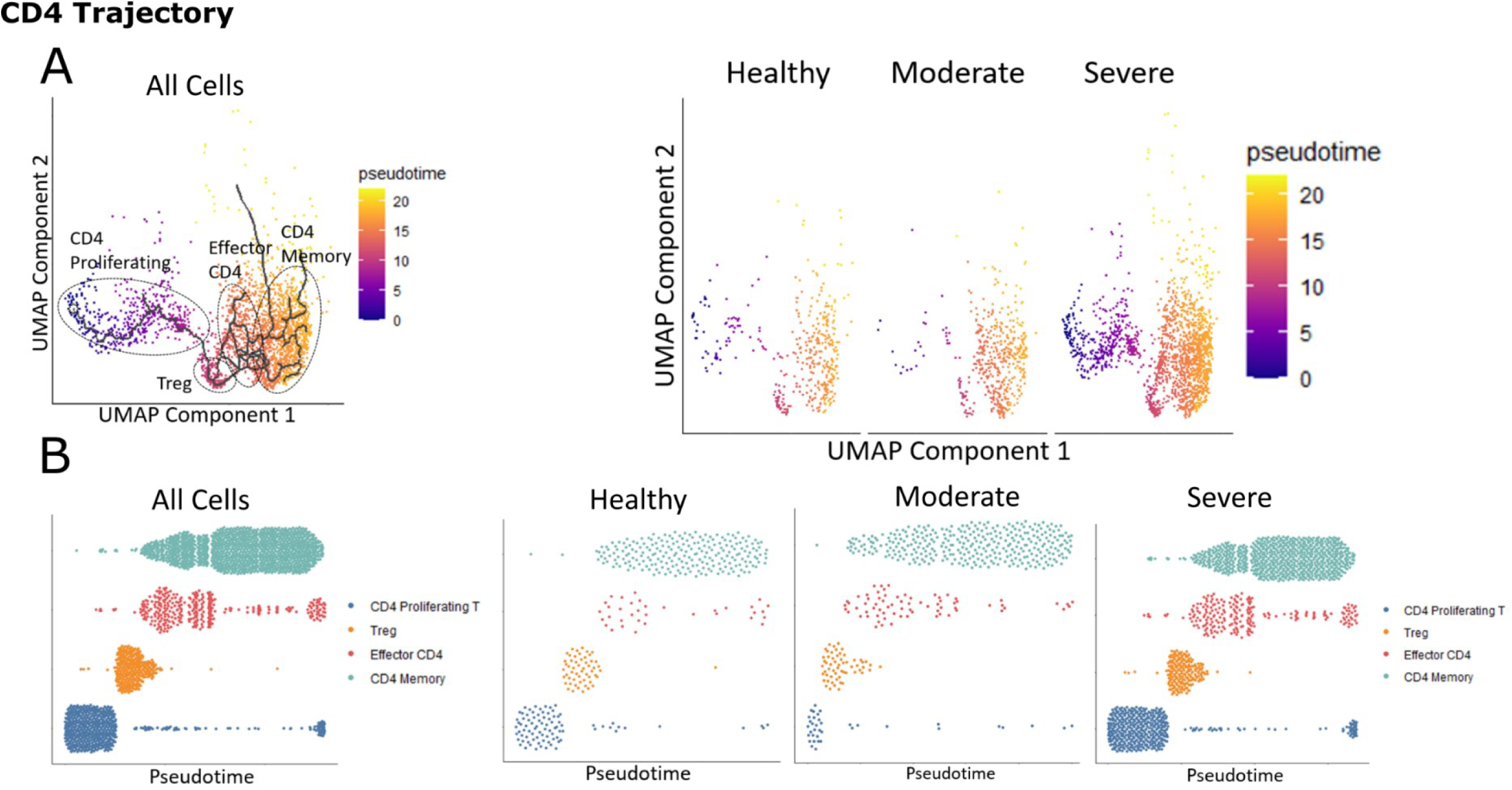
**A**. UMAP projection of 2603 CD4 cells from all reintegrated samples, healthy samples alone, moderate samples alone, and severe samples alone, with trajectory mappings colored by pseudotime; **B**. Dot plot showing pseudotime values for CD4 cells from all reintegrated samples, healthy samples alone, moderate samples alone, and severe samples alone, each dot represents a cell.

**Fig. S8.**
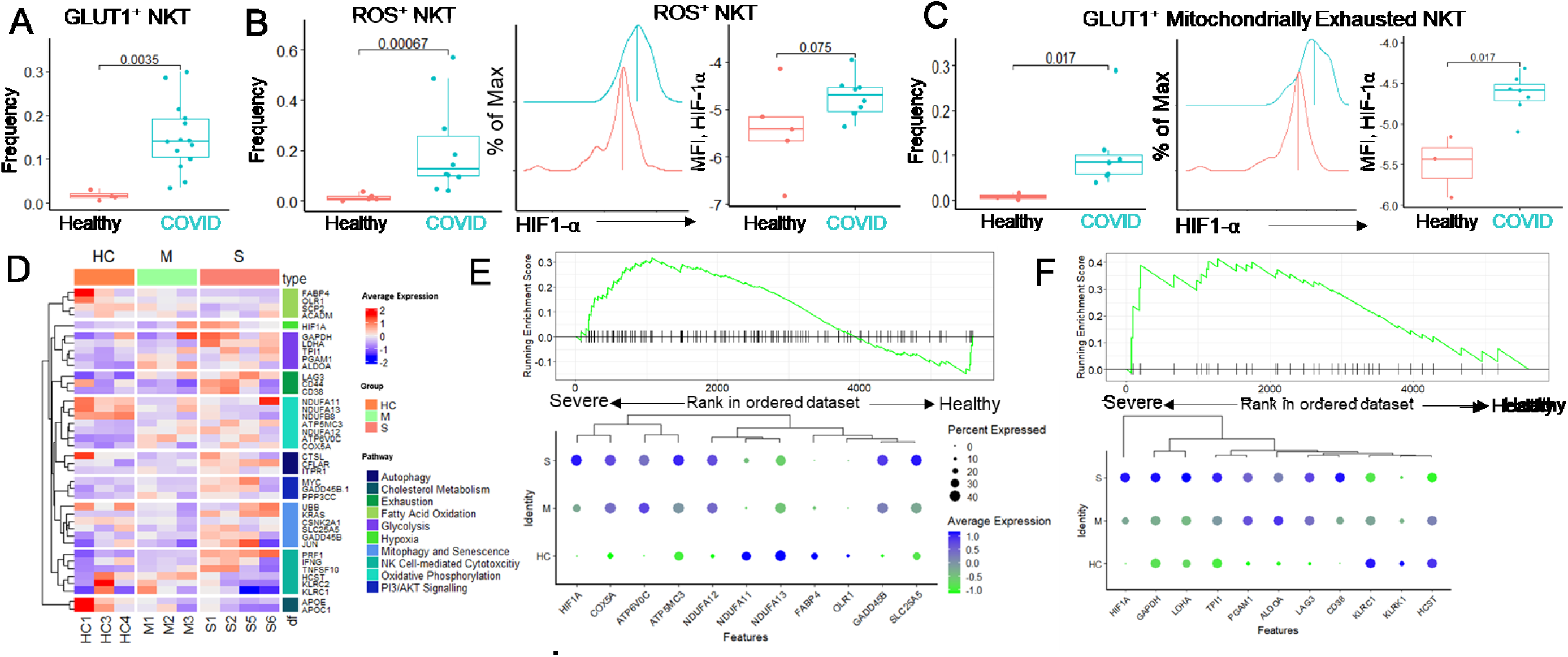
Metabolic profiling of NKT in PBMCs and BALF of COVID-19 patients. NKT cells detected by unsupervised clustering in PBMCs were assessed for expression of metabolic markers and cell function. **A.** Box-plot of cell-type proportion of GLUT1+ NKT cells for each disease state, each dot represents individual sample, 2-sided Wilcoxon Mann Whitney test was performed to indicate statistical significance; **B.** Box-plot of cell-type proportion of ROS+ NKT cells for each disease state, adjacent is density plot displaying distribution of HIF-1a expression in ROS+ NKT cells, alongside boxplot displaying MFI values of HIF-1a expression in ROS+ NKT cells for both Healthy and COVID-19 patients; **C.** Box-plot of cell-type proportion of GLUT1+ Mitochondrially Exhausted NKT cells for each disease state, adjacent is density plot displaying distribution of HIF-1a expression in GLUT1+ Mitochondrially Exhausted cells, alongside boxplot displaying MFI values of HIF-1a expression in GLUT1+ Mitochondrially Exhausted cells for both Healthy and COVID-19 patients; **D.** Heatmap displaying expression of key differentially expressed metabolic genes for NKT cells; **E.** GSEA enrichment plots for “TCA and Respiratory Electron Transport” pathways comparing severe vs healthy control patients, adjacent is a dotplot demonstrating expression and hierarchical clustering of select key genes involved in mitochondrial metabolism; **F.** GSEA enrichment plots for “Glycolysis” comparing severe vs healthy control patients, adjacent is a dotplot demonstrating expression and hierarchical clustering of select key glycolytic genes; **G.** UMAP projections of NKT cells clustered solely on the expression of key differentially expressed metabolic genes; **H.** Correlation matrix showing pearson correlation between differentially expressed metabolic genes

**Supplementary Table 1.**
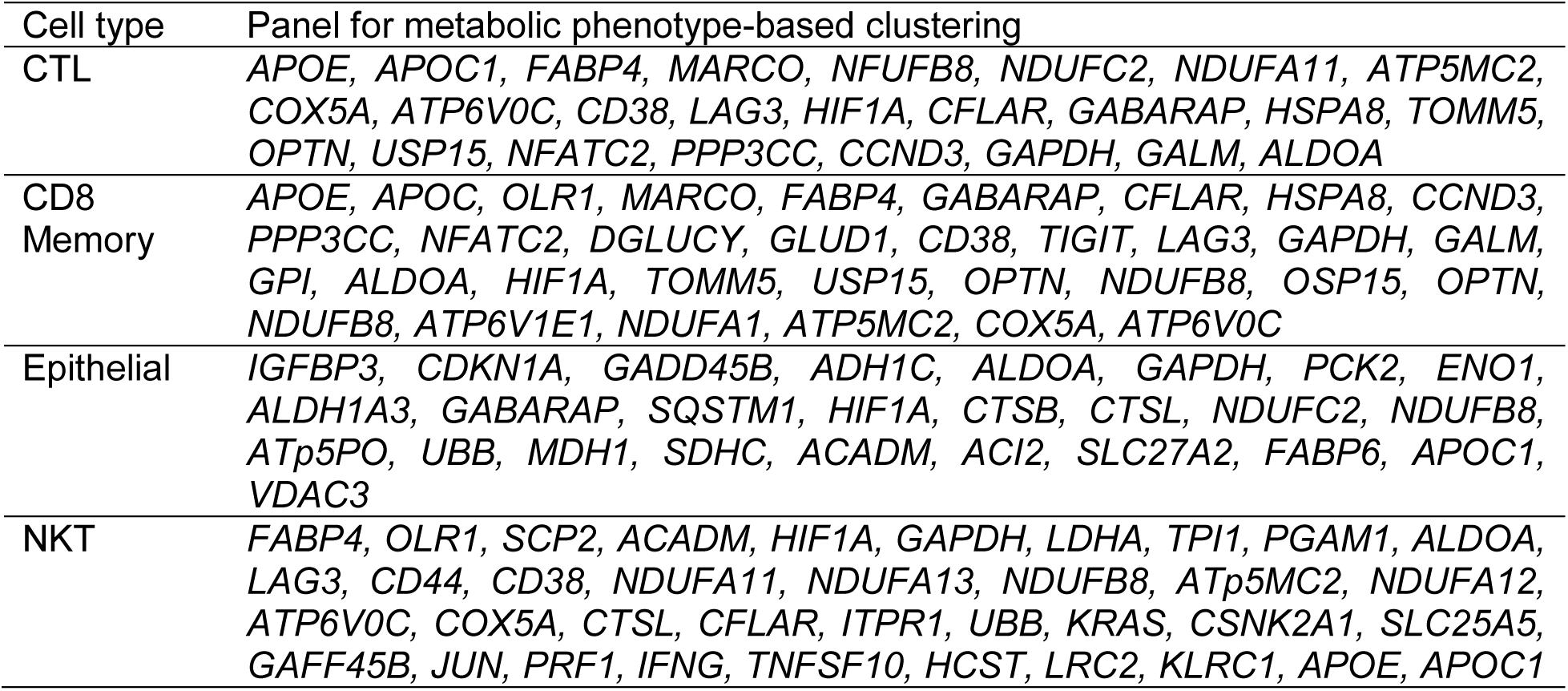

**Supplementary Table 2.**

|  | Glycolysis Module Score | FAO Module Score | HLA Class 2 Signaling Module Score | Type 1 Interferon Module Score | NfKB Module Score |
| --- | --- | --- | --- | --- | --- |
| CD8 Memory | *GAPDH, GALM, GPI, ALDOA* | *OLR, MARCO, FABP4* |  |  |  |
| Epithelial | *ADH1C, ALDOA, GAPDH, PCK2, ENO1, ALDH1A3* | *ACADM, ECI2, SLC27A2, FABP6* | *HLA-DRA, HLA-DPA1, HLA-DMA, DYNLL1* | *IFITM1, IFITM2, IFIT1, MX1, IFITM2* | *SQSTM1, GADD45B, NFKBIA, RELB* |

**Supplementary Table 3.**

| PBMC Cell Population | Surface Markers Used for Identification |
| --- | --- |
| Cytotoxic T Lymphocytes | CD8, GZMB |
| CD8 Central Memory Cells | CD8, CD62L, and CCR7 |
| CD8 Transitional Memory Cells | CD8, CD62L |
| CD8 Effector Memory Cells | CD8, CCR7, GZMB |
| NK Cells | CD56 |
| NKT Cells | CD56, CD8 |
| CD62L<sup>+</sup> NK Cells | CD56, CD62L |

**Supplementary Table 4.**
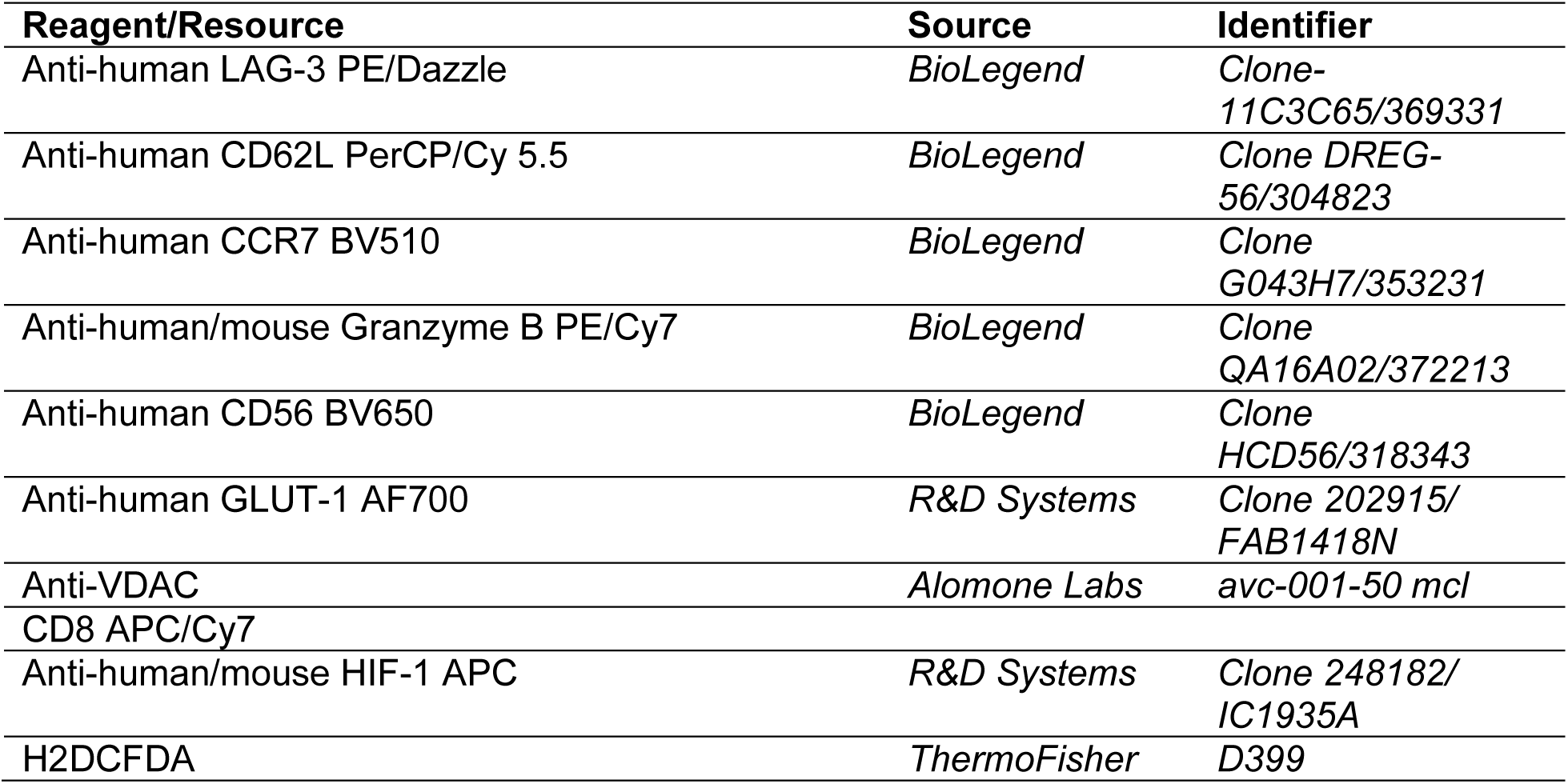

